# Characterization of the genetic architecture underlying eye size variation within *Drosophila melanogaster* and *Drosophila simulans*

**DOI:** 10.1101/555698

**Authors:** Pedro Gaspar, Saad Arif, Lauren Sumner-Rooney, Maike Kittelmann, Andrew J. Bodey, David L. Stern, Maria D. S. Nunes, Alistair P. McGregor

**Affiliations:** Department of Biological and Medical Sciences, Oxford Brookes University, Gipsy Lane, Oxford, OX3 0BP, UK; Centre for Functional Genomics, Oxford Brookes University, Oxford, OX3 0BP, UK; Oxford University Museum of Natural History, Parks Road, Oxford OX1 3PW, UK; Diamond Light Source, Harwell Science & Innovation Campus, Oxfordshire, OX11 ODE, UK; Janelia Research Campus, Howard Hughes Medical Institute, Ashburn, VA 20147, USA

**Keywords:** *Drosophila*, evolution, development, eye size, ommatidia

## Abstract

The compound eyes of insects exhibit striking variation in size, reflecting adaptation to different lifestyles and habitats. However, the genetic and developmental bases of variation in insect eye size is poorly understood, which limits our understanding of how these important morphological differences evolve. To address this, we further explored natural variation in eye size within and between four species of the *Drosophila melanogaster* species subgroup. We found extensive variation in eye size among these species, and flies with larger eyes generally had a shorter inter-ocular distance and *vice versa*. We then carried out quantitative trait loci (QTL) mapping of intra-specific variation in eye size and inter-ocular distance in both *D. melanogaster* and *D. simulans.* This revealed that different genomic regions underlie variation in eye size and inter-ocular distance in both species, which we corroborated by introgression mapping in *D. simulans*. This suggests that although there is a trade-off between eye size and inter-ocular distance, variation in these two traits is likely to be caused by different genes and so can be genetically decoupled. Finally, although we detected QTL for intra-specific variation in eye size at similar positions in *D. melanogaster* and *D. simulans*, we observed differences in eye fate commitment between strains of these two species. This indicates that different developmental mechanisms and therefore, most likely, different genes contribute to eye size variation in these species. Taken together with the results of previous studies, our findings suggest that the gene regulatory network that specifies eye size has evolved at multiple genetic nodes to give rise to natural variation in this trait within and among species.

## Introduction

Animal eyes show striking morphological variation reflecting adaptations to particular habitats, and differences in lifestyle and behaviour (LAND and NILSSON 2012). Insect compound eyes are made up of subunits called ommatidia, which vary in number, structure, and size within and among species causing differences in overall eye size and shape (LAND and NILSSON 2012). Variation in ommatidia number and/or size have important implications for vision: higher numbers of narrower ommatidia can increase acuity, while wider ommatidia can improve contrast sensitivity but increase inter-ommatidial angles, consequently reducing acuity (LAND 1997; LAND and NILSSON 2012). Therefore, optimization of acuity and contrast sensitivity involves trade-offs between ommatidia number and their size, which are determined during eye development.

Compound eye development is understood in great detail in *Drosophila melanogaster* (WOLFF and READY 1993; DOMINGUEZ and CASARES 2005; KUMAR 2011; KUMAR 2012; CASARES and ALMUDI 2016; KUMAR 2018). In this fly, the eyes as well as the head capsule tissue, ocelli, antennae and maxillary palps develop from the eye-antennal disc. Ommatidia number is determined during the third larval instar as the morphogenetic furrow moves from posterior to anterior across the retinal field to trigger the differentiation of rows of ommatidia in their hexagonal pattern (reviewed in FRANKFORT and MARDON 2002; KUMAR 2011; KUMAR 2012). The final size of ommatidia is determined later during the pupal stage as these facets become fully differentiated, although much less is known about the specification of ommatidia size than ommatidia number (CAGAN and READY 1989; VANDENDRIES *et al.* 1996; MILLER and CAGAN 1998; FICHELSON *et al.* 2012; KIM *et al.* 2016).

*D. melanogaster* and other species in this subgroup exhibit substantial variation in eye size caused by differences in the diameter and/or number of ommatidia (NORRY *et al.* 2000; HAMMERLE and FERRUS 2003; POSNIEN *et al.* 2012; ARIF *et al.* 2013; HILBRANT *et al.* 2014; KEESEY *et al.* 2019; RAMAEKERS *et al.* 2019). Like other drosophilids and other dipterans, this variation in eye size is negatively correlated with face width (inter-ocular distance) and/or antennal size, suggesting trade-offs during the development of eye-antennal tissues that contribute to their final size (NORRY *et al.* 2000; DOMINGUEZ and CASARES 2005; SUKONTASON *et al.* 2008; POSNIEN *et al.* 2012; KEESEY *et al.* 2019; RAMAEKERS *et al.* 2019).

Studying the genetic basis of eye size variation in *Drosophila* offers an excellent opportunity to better understand the regulation and evolution of the development of eyes and other tissues, and ultimately how these morphological changes can cause functional differences in vision. For example, strains of *D. mauritiana* have larger eyes than either *D. melanogaster* or *D. simulans*, caused mainly by differences in ommatidial diameter, which is wider in *D. mauritiana* (POSNIEN *et al.* 2012; ARIF *et al.* 2013). QTL mapping has shown that while the larger eyes of *D. mauritiana* is caused by an X-linked locus of large effect, the reciprocal shorter inter-ocular distance of this species is caused by non overlapping autosomal loci (POSNIEN *et al.* 2012; ARIF *et al.* 2013). These effects were verified by introgressing these regions from *D. mauritiana* into *D. simulans* to generate flies with larger eyes or a shorter inter-ocular distance (POSNIEN *et al.* 2012; ARIF *et al.* 2013). Additionally, a genome-wide association study in *D. melanogaster* found an association between variation at the *kek1* locus and female inter-ocular distance but not eye size (VONESCH *et al.* 2016). In contrast, analysis of *D. melanogaster* recombinant inbred lines suggested a common genetic basis for eye and head capsule variation (NORRY and GOMEZ 2017). Furthermore, a polymorphism in the 3^rd^ intron of *eyeless* (*ey*) has recently been shown to contribute to variation in eye size, caused by ommatidia number differences, and reciprocal changes in antennal size/inter-ocular distance between *D. melanogaster* strains (RAMAEKERS *et al.* 2019).

Taken together, these studies suggest that different genetic mechanisms can cause changes in ommatidia size and/or number, and consequently natural variation in the overall size of *Drosophila* compound eyes and the trade-off with inter-ocular distance. In this study we further explore variation in eye and head capsule morphology in *D. melanogaster* and *D. simulans*, and investigate and compare the genetic and developmental bases for intra-specific differences in eye size in these species. This provides new insights into the genetic and developmental bases of eye size variation within and between species.

## Materials and Methods

### Fly strains and husbandry

Multiple strains of *D. melanogaster*, *D. simulans*, *D. mauritiana* and *D. sechellia* were used in this study, including strains from the ancestral range and other populations of *D. melanogaster* and *D. simulans* (Table S1). Flies were maintained on a standard cornmeal diet at 25°C under a 12:12 hour dark/light cycle. For experiments, flies were reared at a low density, achieved through mating of no more than five females and allowing a 5 to 8 hour egg laying period, typically resulting in fewer than 30 larvae per vial.

**Table 1.** *D. melanogaster* QTL for eye area, inter-ocular distance and tibia length

| Trait | QTL<br>(Chr:Mb) | Peak<br>Significance<br>(LOD) | 2LOD Support Region<sup>1</sup> |  |  | QTL Effects |  |  |
| --- | --- | --- | --- | --- | --- | --- | --- | --- |
|  |  |  | Range<br>(Mb) | Start (bp) | End (bb) | Standardized<br>Additive<br>Effect<sup>2</sup> | Standardized<br>Dominant<br>Effect<sup>3</sup> | Variance<br>Explained (%) |
| Eye area | 3L:4.51 | 2.487 | 8.35 | 3L:<br>4512922 | 3R:<br>18043455 | 0.20 | 0.06 | 3.931 |
| Inter-ocular<br>distance | 2R:24.74 | 4.251 | 47.60 | 2L:<br>661886 | 2R:<br>24746955 | 0.31 | -0.11 | 6.322 |
|  | 3L:0.82 | 3.939 | 2.080 | 3L:<br>821255 | 3L:<br>10527569 | 0.11 | 0.13 | 3.011 |
| Tibia 2<br>length | 2L:9.01 | 5.553 | 10.35 | 2L:<br>661886 | 2L:<br>21292443 | -0.37 | 0.10 | 5.905 |
|  | 3L:0.82 | 1.452 | 41.53 | 3L:<br>821255 | 3R:<br>153094825 | 0.04 | 0.13 | 1.467 |
|  | 3R:15.30 | 2.628 | 22.58 | 3L:<br>6019468 | 3R:<br>29777199 | -0.13 | 0.25 | 2.694 |
| Tibia 3<br>length | 2L:1.63 | 2.203 |  | 2L:<br>661886 | 2R:<br>24746955 | -0.23 | 0.02 | 2.638 |
|  | 3L:6.01 | 5.051 | 50.34 | 3L:<br>821255 | 3R:<br>21255695 | -0.12 | 0.36 | 6.268 |

<sup>1</sup>Regions corresponding to the 2-LOD support interval based on the next marker position closest to the interval boundaries.
<sup>2, 3</sup>See Materials and Methods for details on how these different measures of effect size were calculated.

### Phenotypic measurements

All parental strains were imaged from frontal or lateral views of the head and the *D. melanogaster* QTL mapping population was imaged from frontal head views, captured by a Axiocam 506 colour D camera mounted on a Zeiss AxioZoom.V16 microscope with an Apo Z 1.5x/0.37 FWD 30 mm objective. For the *D. simulans* mapping population, frontal images of the head were taken using a Leica M205 stereomicroscope and a DFC300 camera. Eye area was measured as the sum of outlined eye taken from frontal images of the head, as previously described (POSNIEN *et al.* 2012), and the width of the cuticle between the eyes (face width/inter-ocular distance) was measured at the height of the orbital bristles just above the antennae. All frontal images were further annotated with a combination of 45 landmarks and semi-landmarks as previously described (POSNIEN *et al.* 2012) (Fig. 2C). The landmarks coordinates were then subjected to a Generalized Procrustes Analysis (GPA) to standardize for size, position and orientation. We analyzed variation in head shape using Principal Component Analysis (PCA) of the GPA aligned configurations of head shapes and visualized these differences using thin-plate spline (TPS) deformation grids. All morphometric analysis was performed using the ‘geomorph’ R package (ADAMS and OTÁROLA-CASTILLO 2013). PC1 corresponded to differences in head posture during image acquisition and hence we discarded this PC. The length of the middle (T2) or the most posterior (T3) leg tibias were measured as a proxy for overall body size. All linear measurements and landmark annotations were performed with the Fiji image analysis software (SCHINDELIN *et al.* 2012). Statistical analysis of species, sex and strain differences in eye area, inter-ocular distance and tibia length within the *D. melanogaster* species subgroup survey was carried out using ANOVA and individual differences were tested using the Tukey’s multiple comparisons test.

### X-ray imaging, reconstruction and analyses

Fly heads were dissected under CO_2_ and placed into PBS and later fixed in Bouin’s solution (75 parts saturated aqueous picric acid, 25 parts formalin, 5 parts acetic acid) (PRESNELL and SCHREIBMAN 1997) for 10 hours, washed in 70% ethanol and left for dehydration in 70% ethanol for 12 hours. Heads were then stained with 1% iodine in 70% ethanol for 14 hours, washed in 70% ethanol and left at 4°C in fresh 70% ethanol until imaging. Fly heads were mounted in 20 µl pipette tips filled with 70% ethanol for synchrotron radiation X-ray tomography (SRXT) and scanned at the Diamond-Manchester Imaging Branchline I13-2 [refs] (Diamond Light Source, UK) (RAU *et al.* 2011; PEŠIĆ *et al.* 2013). A partially-coherent, near-parallel, polychromatic ‘pink’ beam was generated by an undulator in an electron storage ring of 3.0 GeV voltage and 280 mA current. The beam was reflected from the platinum stripe of a grazing-incidence focusing mirror and high-pass filtered with 1.3 mm pyrolytic graphite and 3.2 mm aluminium, resulting in a weighted-mean photon energy of ∼27 keV. The propagation (sample-to-scintillator) distance was set to ∼50mm, to give a low level of inline phase contrast. The undulator gap was set to 5 mm for data collection and 10 mm for sample alignment. Effective beam area was restricted to ∼1.1 × 1.0 mm for data collection; this limited both sample exposure and the intensity of noise arising from scintillator defects. 4001 projection images were acquired at equally-spaced angles over 180° of continuous rotation (‘fly scan’), with an extra projection (not used for reconstructions) collected at 180° to check for possible sample deformations, bulk movements and beam damage relative to the first (0°) image. Images, of 140 ms exposure time, were collected by a pco.edge 5.5 (PCO AG, Germany) detector (sCMOS sensor of 2560 × 2160 pixels) mounted on a visual light microscope of variable magnification. A 10x objective, coupled to a GGG:Eu scintillator and mounted ahead of a 2x lens, provided 20x total magnification, a field of view of 0.8 × 0.7 mm and an effective pixel size of 325 nm, as confirmed with a laminographic standard. Data were reconstructed using a filtered back projection algorithm in the modular pipeline Savu 2.3 (ATWOOD *et al.* 2015; WADESON and BASHAM 2016), incorporating flat- and dark-field correction, optical distortion correction (VO *et al.* 2015; STROTTON *et al.* 2018), ring artefact suppression (KIM *et al.* 2014) and Paganin filtering (δ/β=4). Images were transformed into 16 bit. The IMOD Software package (KREMER *et al.* 1996) was used to generate mrc stacks from reconstructed tomogram TIFF files. Stacks were binned 2x in X and Y to reduce file size for 3D segmentation in Amira v.2019.2 (Thermo Fisher Scientific). Ommatidial facets and head structures were segmented generating a mask material around the cornea and head cuticle respectively, and then applying the threshold tool within that mask to segment the facets or head. Ommatidial diameter was measured with the 3D line measurement tool on the segmented eye from the dorsal to ventral side of the facets. Differences in ommatidia number and diameter differences were assessed using t-tests.

### Genotyping

DNA was extracted from adult fly abdomens. Genotyping for the *D. melanogaster* QTL mapping population was performed with restriction fragment length polymorphisms (RFLP) at 42 loci regularly spaced across all four chromosomes (Tables S2, S3). The mean distance between consecutive markers was 6.64 cM, with the maximum distance between markers was 15.3 cM. *D. melanogaster* genomes were obtained from the Drosophila Genome Nexus and aligned to the *D. melanogaster* reference genome BDGP release 5 (POOL *et al.* 2012) (SRP005599, http://www.johnpool.net/genomes.html). Genotyping for the *D. simulans* mapping population was performed using multiplexed shotgun genotyping (MSG) (ANDOLFATTO *et al.* 2011) with 6152 SNPs for the backcross to the Tana10 strain and 8115 SNPs for the backcross to the Zom4 strain (Tables S4-S9). Parental genomes were generated by updating the *D. simulans* r2.0.1 genome (http://www.flybase.org/static_pages/feature/previous/articles/2015_02/Dsim_r2.01.html) with HiSeq reads from each strain (these genomes are available on request). Genotypes were estimated using the MSG software (https://github.com/YourePrettyGood/msg/tree/dev).

**Table 2.**
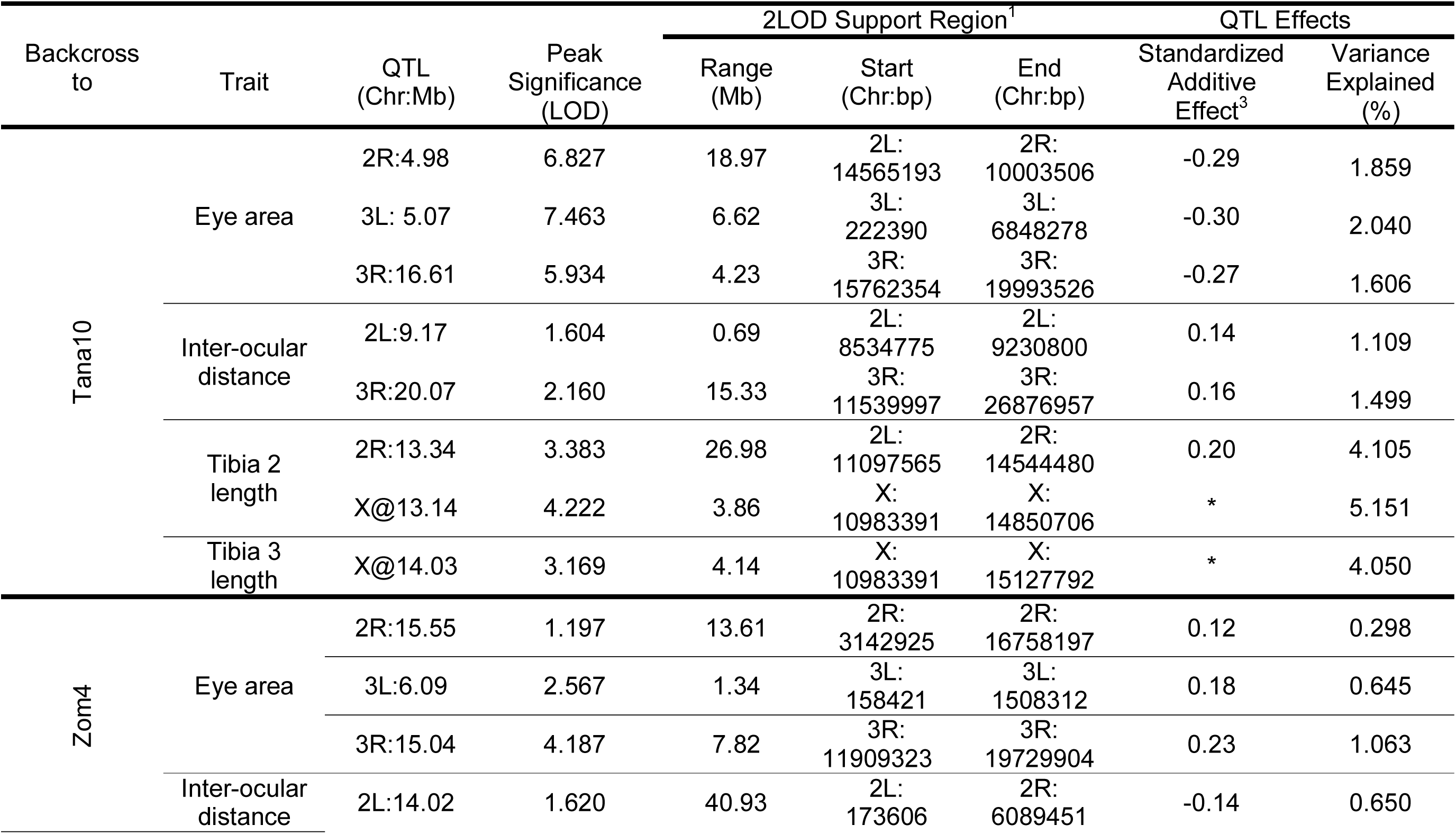
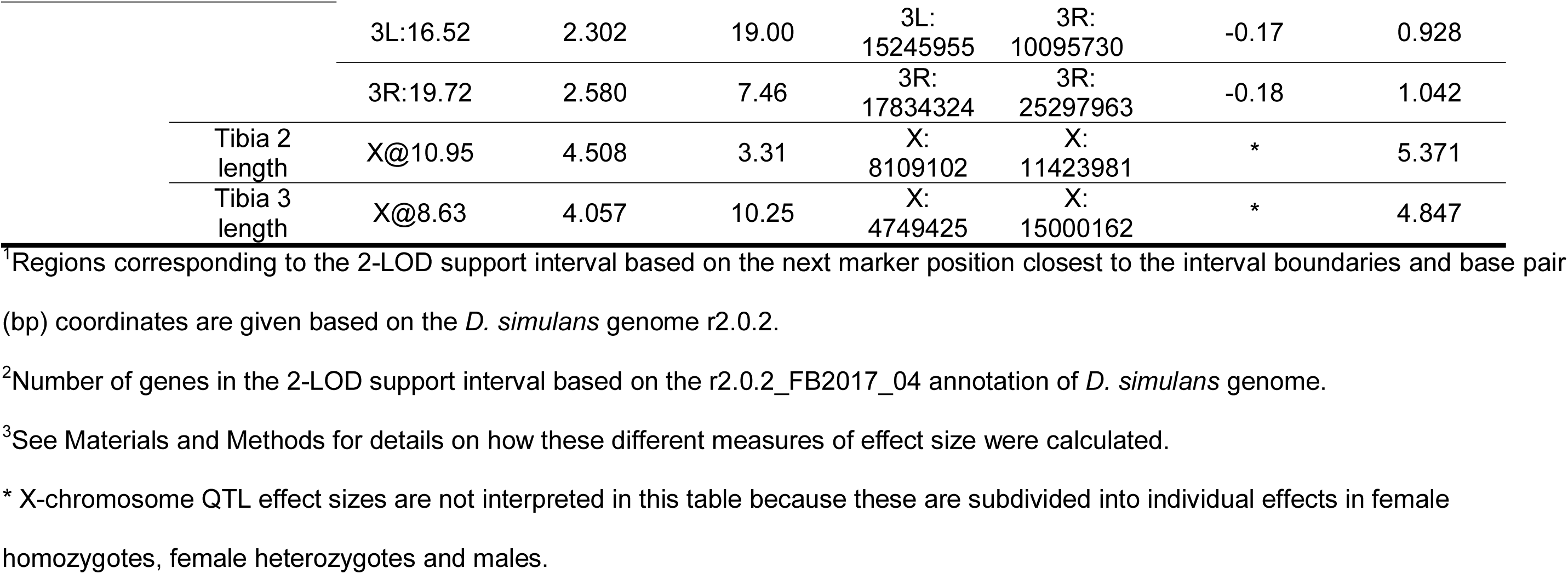
*D. simulans* QTL for eye area, inter-ocular distance and tibia length

### QTL mapping and statistical analyses

To generate the *D. melanogaster* QTL mapping population, *D. melanogaster* ZI373 females were mated to *D. melanogaster* RG13N males, or reciprocally (Fig. S1A). F1 progeny from each reciprocal parental cross were mated to siblings as density-controlled replicates of 5 females crossed to 5 males. The heads of 48 F2 females and 48 F2 males from each cross direction (totaling 192 individuals) were phenotyped and their bodies processed for genotyping with RFLPs (Table S3, Fig. S1A). To generate the *D. simulans* QTL mapping population, *D. simulans* Zom4 females were mated to *D. simulans* Tana10 males. F1 virgin females were then backcrossed to either Zom4 males or Tana10 males as density controlled replicates of 5 females crossed to 5 males (Fig. S1B). The heads of 192 female and 192 male progeny from each backcross were phenotyped and their bodies were processed into one MSG library per backcross (Tables S4-S9, Fig. S1B). For the *D. simulans* genotype dataset, posterior probabilities of ancestry were thinned and imported into R/qtl using custom scripts (http://www.github.com/dstern/pull_thin, http://www.github.com/dstern/read_cross_msg).

To determine QTL locations for both the *D. melanogaster* and *D. simulans* crosses, we performed genome scans with a single QTL model using R/qtl (as implemented by the ‘scanone’ function) to perform standard interval mapping with Haley-Knott regression (HALEY and KNOTT 1992; BROMAN *et al.* 2003). Genome**-**wide statistical significance thresholds (0.05%) were determined for each phenotype using 1000 permutations. For both the *D. melanogaster* and *D. simulans* QTL analyses we filtered any individuals with > 10% missing data and any markers with > 10% missing data. Additionally, for *D. simulans*, we only retained markers that were at least 2.5 kb apart for computational efficiency. To identify QTLs in *D. melanogaster* and *D. simulans* we used the native R/qtl forward search/backward elimination search algorithm (as implemented using the ‘stepwiseqtl’ function) followed by a final scan for any additional QTL, after accounting for those discovered in the previous step. We used the lengths of T2 and T3 tibias (as proxies for body size) and sex as covariates, to account for effects of body size and sexual dimorphism, while searching for QTLs associated with eye size and inter-ocular distance. Only sex was used a covariate while searching for QTL associated with tibia lengths. We calculated 2 LOD support intervals for all significant QTL and tested for pair wise interactions between all significant QTL by fitting full linear models in an ANOVA framework (F tests, type III sum of squares), with all significant QTL and a proxy for body size as fixed effects. Furthermore, we estimated additive allelic effects of all significant QTL in three ways for autosomes: (1) the additive effect as half the standardized difference between the means of the homozygotes (ZI373/ZI373, RG13N/RG13N, Zom4/Zom4, Tana10/Tana10), (2) the dominant effect (only for the F2 cross) as the standardized difference between the mean of the heterozygotes (ZI373/RG13N) and the average of the homozygote means (and ZI373/ZI373, RG13N/RG13N), (3) the percentage of phenotypic variance accounted for by the significant QTL in the mapping population (variance explained).

### Introgression design

To generate *D. simulans* introgression lines, *D. simulans* Zom4 females were mated to *D. simulans* Tana10 males, or reciprocally. F1 male progeny were then backcrossed to the maternal line and individual backcross male progeny were again backcrossed to the maternal line (Fig. S2A). These backcross males were genotyped after mating to select for lines that inherited a single autosome from the paternal line, based on RFLPs (Fig. S2A, Table S10). The progeny of such lines were then crossed with each other, as sibling pairs. Each sibling pair was genotyped after mating to select for heterozygous parents for the introgressed autosome of interest. The progeny of such sibling pairs were again sib-pair mated and these were genotyped after mating at multiple RFLPs to selected for homozygous parents for the introgressed autosome of interest (Fig. S2A, Table S10). Whole chromosome introgressions were fixed at this point, or further sib-pair matings were carried out to isolate partial chromosome introgressions with one or more recombination breakpoints. The same strategy was used to generate *D. melanogaster* chromosome introgression lines between strains ZI373 and RG13N (Fig. S2A, Table S2).

Higher resolution introgression mapping in *D. simulans* made use of the whole chromosome introgression of the Zom4 chromosome 3 into the Tana10 background, crossed to the Tana10 strain containing a single Pax3>dsRed transgene inserted at either the left tip (*Dsim*r2.02 3L:761452) or the right tip (*Dsim*r2.02 3R:25786766) of chromosome 3 (Fig. S2B). F1 female progeny of the above cross were backcrossed to the whole-chromosome introgression line. Backcross male progeny were anesthetized and screened using a fluorescent Zeiss AxioZoom.V16 microscope and selected based on inheritance of the Pax3>EGFP or Pax3>dsRed transgenes, visualized as fluorescent markers in adult eyes. These males were again backcrossed individually to the whole chromosome introgression line and genotyped after mating to identify recombinant lines between the marked Pax3>EGFP or Pax3>dsRed chromosome and the introgressed chromosome, using RFLPs (Fig.S2B, Table S10). Lines carrying specific recombination breakpoints were selected for further sib-pair crosses to generate homozygous introgression lines as described above (Fig. S2B, D).

### CRISPR/Cas9 mutagenesis

To generate marker insertions of the Pax3>dsRed transgene for *D. simulans* Tana10, we first generated introgression lines carrying a X chromosome bearing a nanos>Cas9 transgene, derived from the *D. simulans* strain w501 (STERN *et al.* 2017). These introgressions were generated through a male backcross scheme as reported for introgression mapping, making use of RFLPs to select lines that carry the introgressed X chromosome in the homozygous genetic background of the Tana10 strain. Embryos from this strain were injected with a mix of pCFD3 plasmid containing a guide RNA targeting either the left (CCGATCTTACCAGCCAGCTGGC) or right (CCTCTTAATGGTCAGCCAGGTGC) ends of the 3^rd^ chromosome, with a pHD-dsRed.attP donor plasmid for homology-mediated repair, containing 1 kb homology arms directly abutting the guide RNA cut sites (Gratz *et al.* 2014; Port *et al.* 2014). Guide RNAs were designed to target intergenic regions devoid of known genes or gene regulatory regions, according to Flybase genome annotation *D.sim* r2.0.2_FB2017_04. After injection, G0 animals were backcrossed to the respective injected strains followed by screening of fluorescent eyes in F1 progeny. Microinjections, G0 crosses and F1 screening was carried out by the Cambridge Fly Facility. Lines carrying a homozygous Pax3>dsRed transgene were isolated from sib-pair crosses after PCR amplification of the transgene cassette from the immediately surrounding genomic regions, as opposed to PCR amplification of the intervening genomic region without the transgene, or both.

### Introgression mapping and statistical analysis

We used linear mixed models to determine significant associations between eye area or inter-ocular distance and 31 marker genotypes produced from RFLPs (Tables S10, S11). First, we constructed a reference model with sex, tibia of the third leg (proxy for body size covariate) and the food batch the individual fly was reared in as fixed effects, and the introgression line genotype as a random effect. For each marker genotype we constructed a second model with an additional term for the marker genotype in question as a fixed effect. P-values for each marker were obtained from a likelihood ratio test between the reference model and the model with the additional marker term. We used a Bonferroni threshold to determine significant association between phenotype and markers. The linear mixed models and likelihood ratio tests were carried out using the ‘lme4’ R package (BATES *et al.* 2015).

### Immunohistochemistry

Imaginal discs were dissected in phosphate buffer saline (PBS) and fixed in 4 % (v/v) formaldehyde in PBS for 30 min. Following three washes with PBS, samples were permeabilised in 0.3% (v/v) Triton X-100 in PBS (PBST), then blocked in 5% normal goat serum (Sigma) in PBST before incubation with primary antibodies in this solution overnight. Secondary antibodies were incubated with samples for 2 hours at 4°C before mounting in 80% (v/v) glycerol in PBS. Primary antibodies used were: mouse anti-Eyes absent (Eya) (1:10, DCAD2, Developmental Studies Hybridoma Bank) and rat anti-Elav (1:200, 7E8A10, DSHB). Secondary antibodies used were goat anti-rat or anti-mouse, Alexa 488 and goat anti-rat or anti-mouse Alexa 647 (Molecular Probes), at 1:500 dilutions. Nuclear staining was performed using DAPI (Roche). Fluorescence images were acquired using a Zeiss LSM880 confocal microscope. Tissue surface area was measured across apical and basal optical sections using the outline tools of the Fiji image analysis software.

## Results

### Variation in eye size within and between Drosophila species

We and others have previously shown that species in the *D. melanogaster* complex exhibit substantial intra- and interspecific variation in eye size and inter-ocular distance (NORRY *et al.* 2000; HAMMERLE and FERRUS 2003; DOMINGUEZ and CASARES 2005; POSNIEN *et al.* 2012; ARIF *et al.* 2013; HILBRANT *et al.* 2014; KEESEY *et al.* 2019; RAMAEKERS *et al.* 2019). To widen the survey and to further explore intra-specific variation, we measured the eye size, inter-ocular distance and tibia lengths of 26 strains of *D. melanogaster* and 13 strains of *D. simulans* from around the world, including from their ancestral range, as well as 4 representative strains of *D. mauritiana* and 3 of *D. sechellia* (Fig. 1, Fig. S3, Table S12). We detected striking differences in eye area and inter-ocular distance between species, sexes and among strains of the same species (Fig. 1A, Fig. S3, Table S13). While the degree of sexual dimorphism in eye size varied widely across strains, we did not detect a significant species by sex interaction for eye area (p = 0.669) or inter-ocular distance (p = 0.357) variation in this survey (Fig. 1A, Fig. S3, Table S13).

**Fig. 1.**
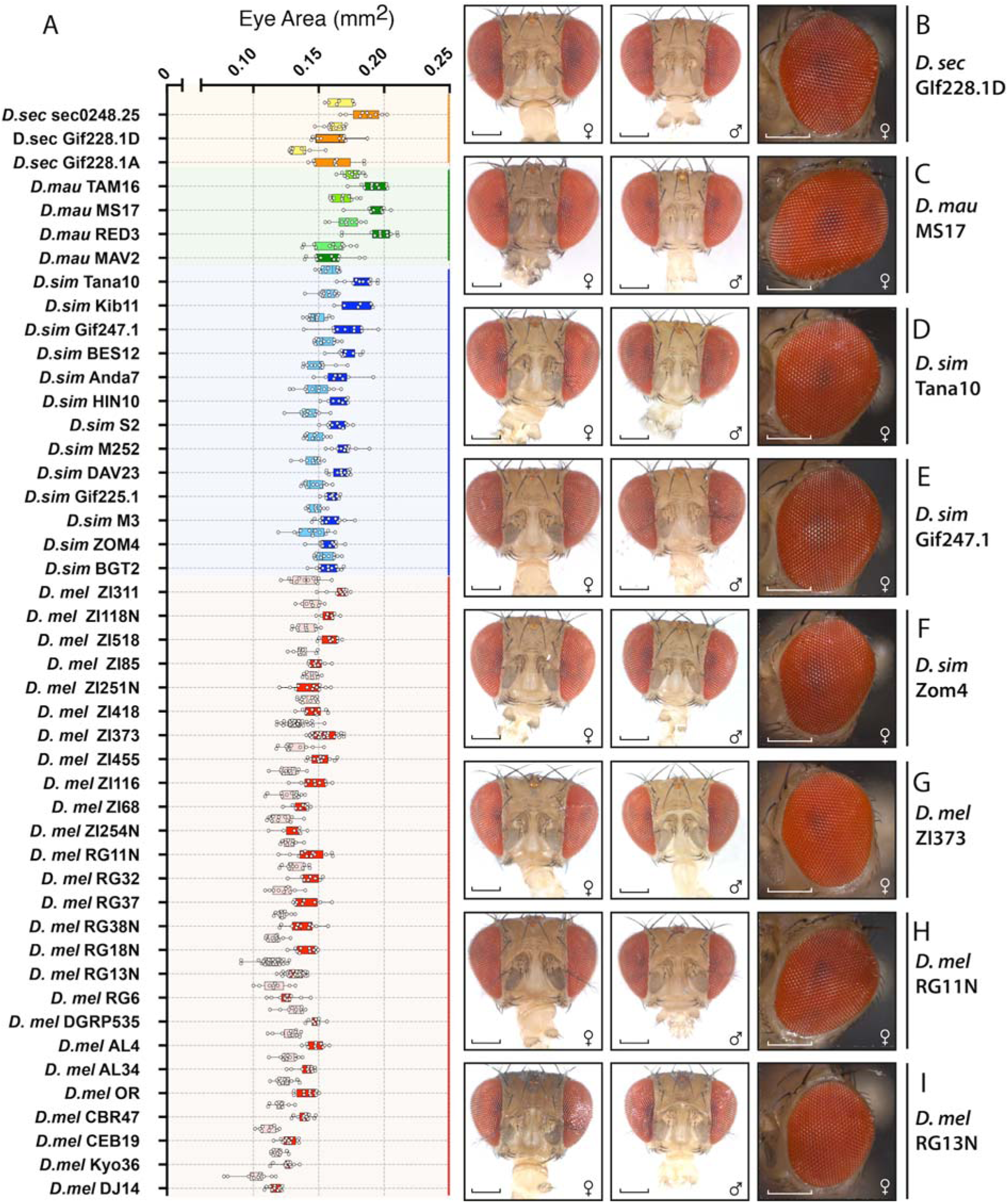
Survey of eye size variation in the *D. melanogaster* species subgroup. (A) Box and whisker diagram of eye area (mm^2^) of strains of *D. sechellia* (males – yellow, females – orange), *D. mauritiana* (males – light green, females – dark green), *D. simulans* (males – light blue, females – dark blue) and *D. melanogaster* (males – pink, females – red). (B, C) Frontal and lateral head views of females (♀) or males (♂) from strains representative of the average of the surveyed eye size variation in *D. sechellia* (B) and *D. mauritiana* (C). (D-I) Frontal and lateral head views of females (♀) or males (♂) from strains representing average and further analysed extreme ends of variation in *D. simulans* (D-F) *and D. melanogaster* (G-I). Scale bar = 200µm (see Table S12 for means, standard deviations and sample sizes and Table S13 for summary statistics).

**Fig. 2.**
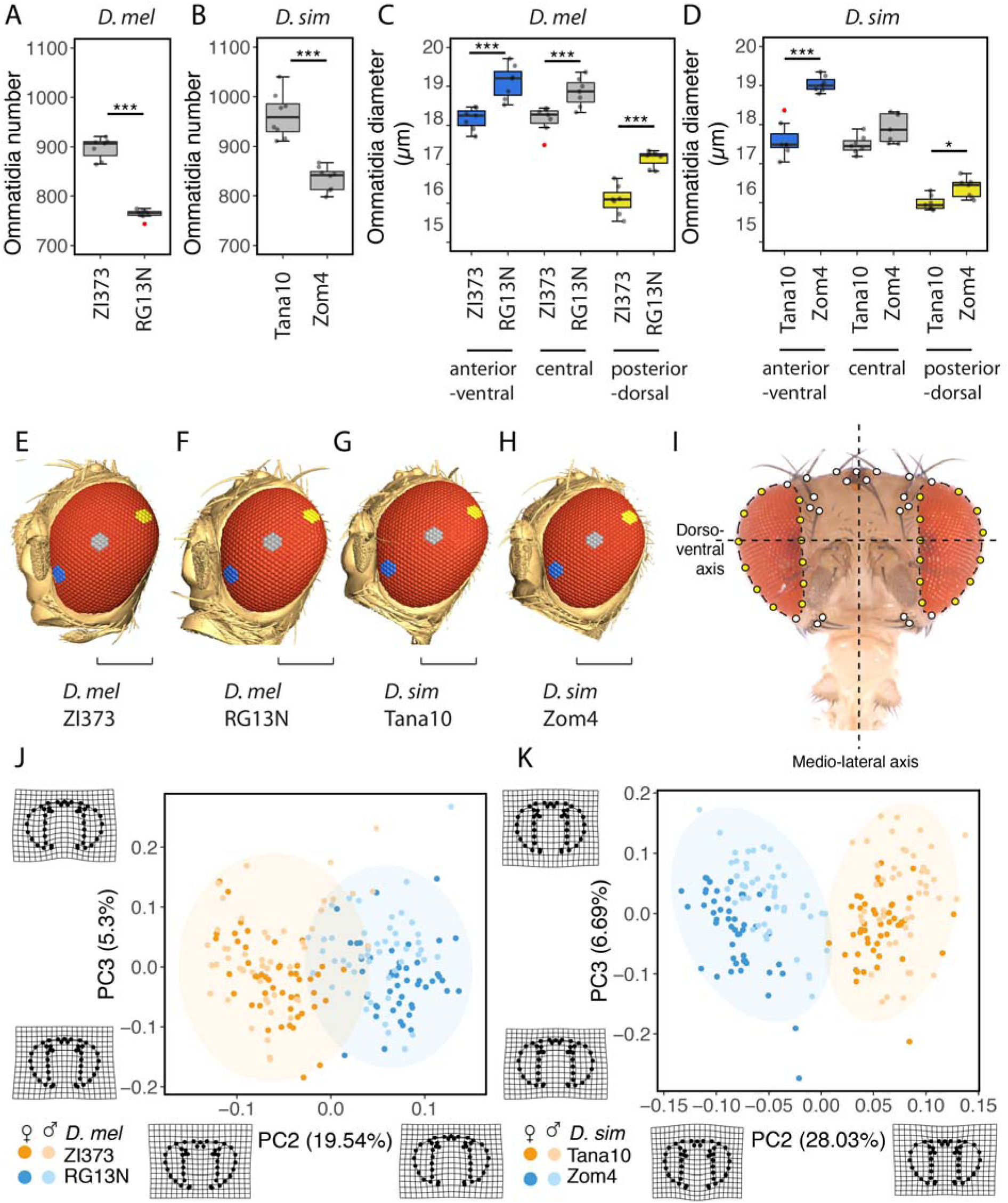
Characterization of the ommatidial bases of eye size and shape differences in focal strains of *D. melanogaster* and *D. simulans*. (A-B) Ommatidia number in females of *D. melanogaster* strains ZI373 (n = 7, µ = 897±22) and RG13N (n = 8, µ = 764±9) (A), and *D. simulans* strains Tana10 (n = 8, µ = 962±45) and Zom4 (n = 8, µ = 835±24) (B). (C,D) Ommatidia diameter (n = 14) measured in anterior-ventral (blue), central (grey) and posterior dorsal (yellow) ommatidia from females of the *D. melanogaster* strains ZI373 (n = 7) and RG13N (n = 7) (C), and *D. simulans* strains Tana10 (n = 7) and Zom4 (n = 7) (D). (E-H) Segmentations from synchrotron X-ray tomograms of female heads of ZI373 (C), RG13N (D), Tana10 (E) and Zom4 (F). Measured ommatidia within anterior-ventral, central and posterior-dorsal groups are highlighted in blue, grey and yellow,, respectively. Scale bar = 200 µm. (I) Position of landmarks (white) and semi-landmarks (yellow) along the front view of a *Drosophila* head used for PCA of head shape. (J,K) Distribution of PC2 and PC3 and their 95% confidence ellipses for position of head landmarks of the *D. melanogaster* strains ZI373 (orange) and RG13N (blue) (J), and *D. simulans* strains Tana10 (orange) and Zom4 (blue) (K). Wireframe deformation diagrams represent 2x magnifications of the minimum and maximum coordinates along the PC axis to illustrate shape differences. In (A-D), red data points represent outliers. Statistical comparisons represent the results of t-tests: *** p<0.0001; *p<0.01.

Our survey showed that *D. melanogaster* females generally have significantly smaller eye areas than those of the other species, being on average 15%, 17% and 24% smaller than *D. simulans*, *D. sechellia* and *D. mauritiana* eyes, respectively (Table S13). However, *D. melanogaster* strains with the largest eyes overlap in eye size with *D. simulans* and *D. sechellia* (Fig. 1, Table S12). Consistent with previous findings, species with large eyes generally exhibit narrow inter-ocular distances and *vice versa* (Fig. 1A, Fig. S3A) (NORRY *et al.* 2000; POSNIEN *et al.* 2012; ARIF *et al.* 2013; KEESEY *et al.* 2019). Interestingly, within-species variation in head traits may reflect population based differences, as observed between the *D. melanogaster* strains from Rwanda (RG) compared to Zambia (ZI), with Rwanda female eyes (0.137±0.0096 mm^2^) being on average 9% smaller than those of Zambia females (0.150±0.0121 mm^2^) (Fig. 1A, Tables S12, S13).

### Analysis of intra-specific variation in D. melanogaster and D. simulans

To study intra-specific variation in eye size in more detail, we focused on pairs of strains of *D. melanogaster* and *D. simulans* with significant differences in eye area: *D. melanogaster* ZI373 versus RG13N and *D. simulans* Tana10 versus *D. simulans* Zom4 (Fig. 1, Fig. S4, Table S14). Between these strains, differences in eye size are inverse to differences in inter-ocular distance: the strain with larger eyes has shorter inter-ocular distance and *vice versa* (Fig. 1, Fig. S4). Furthermore, differences in eye size are consistent with differences in ommatidia number, as females of the ‘large eye’ strains ZI373 and Tana10 have on average 133 and 127 more ommatidia than those of the ‘smaller eye’ strains RG13N and Zom4, respectively (Fig. 2A,B, Table S14). In contrast, ommatidia diameters are significantly wider across the eye in females of the ‘smaller eye’ strains (Fig. 2C-H, Table S14). This difference is particularly noticeable for the anterior-ventral ommatidia between the two *D. simulans* strains, while the *D. melanogaster* strains differ significantly in ommatidia size in all three eye regions studied (Fig. 2C,D, Table S14). Overall, however, strains with smaller ommatidia nevertheless have larger eyes and so the differences in eye size between the focal *D. melanogaster* and *D. simulans* strains can be mostly attributed to the differences in ommatidia number.

To further study differences in the eye and head capsule of the analyzed strains, we used PCA of the position of stereotypical landmarks on frontal views of the heads (Fig. 2I). Thin-plate splines interpolation illustrates head shape differences along the medial-lateral axis (Fig. 2J,K), consistent with the inverse relationship between eye size and inter-ocular distance between strains. Interestingly, differences along the dorsal-ventral head axis indicate that the eyes of larger eyed strains ZI373 and Tana10 are relatively wider in the dorsal region compared to the smaller eyed strains RG13N and Zom4, which appear more symmetric along this axis (Fig. 2J,K).

Eye size differences between strains could reflect differences in overall body size, as eye area is generally positively correlated with tibia length in the strains of both species (Figs S5). Indeed, for *D. simulans*, Tana10 females have significantly longer tibias than those of Zom4 (Fig. S4E). However, residuals of regression of eye area with tibia length show that the eyes of Tana10 are still significantly larger than Zom4 independently of this proxy for body size (Fig. 3A). In *D. melanogaster*, the strain with larger eyes ZI373 has shorter tibias compared to the strain with smaller eyes RG13N (Fig. S4C). When comparing the residuals of regression of eye area on tibia length, ZI373 eyes are still significantly larger than those of RG13N, but comparing the residuals for inter-ocular distance with tibia length between ZI373 and RG13N, shows that ZI373 has narrower inter-ocular distance after this correction (Fig. S6A). This illustrates that there are differences in the scaling relationships between tibia length and eye area or inter-ocular distance. Therefore, when further studying eye size and inter-ocular distance below, we considered both traits with and without correcting for tibia length (Fig S6A).

**Fig. 3.**
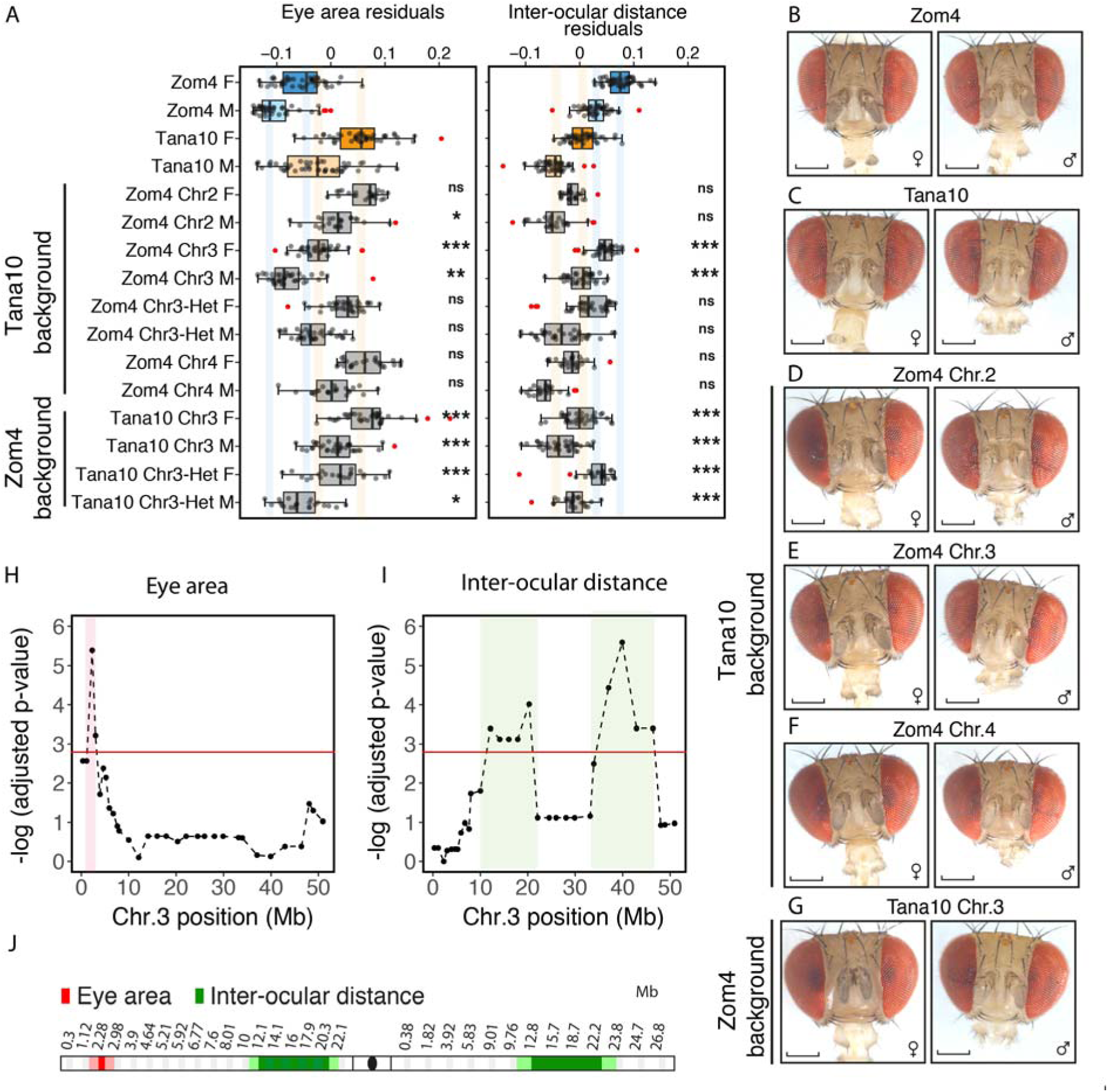
Genetic mapping of eye size and inter-ocular distance variation using *D. simulans* introgressions. (A) Box and whisker diagram of eye area (mm^2^) and inter-ocular distance (µm) measured for both females (F) and males (M) of the parental strains Zom4 and Tana10 and introgression of chromosome 2 (Chr2), chromosome 3 (Chr3) and chromosome 4 (Chr4) from these strains into either genetic background. (B-G) Frontal head views of females (♀) or males (♂) from strains representative of the average of the parental strains (B) Zom4 and Tana10 (C) comparing with introgression of chromosome 2 (D), 3 (E) and 4 (F) of Zom4 into the Tana10 background and of chromosome 3 of Tana10 into the Zom4 background (G). Log-adjusted p-values for a linear mixed model comparing fixed versus random association of eye area (H) or inter-ocular distance (I) with marker genotypes along the 3^rd^ chromosome for partial introgressions of Tana10 or Zom4 chromosome 3 into either strain genetic background (see Fig.S2D for details of recombination breakpoints and genetic background). (J) Candidate regions along chromosome 3 positions (Mb) are highlighted in red for eye area and in green for inter-ocular distance. scale bar – 200 µm. Statistical comparisons represent pairwise t-tests by sex between parental strains (red asterisks) or between introgression line and the parental strain by sex to which this line was backcrossed to (black asterisk): ***p<0.0001, **p<0.001, *p<0.01, ns – p>0.01.

### *QTL mapping in* D. melanogaster and D. simulans

To investigate and compare the genetic basis of intra-specific differences in eye size and inter-ocular distance in *D. melanogaster* and *D. simulans* we carried out QTL mapping on these traits in both species with and without sex and tibia length as co-variates.

In *D. melanogaster* we generated a mapping population of 192 F2 individuals from reciprocal crosses between strains ZI373 and RG13N (Fig. S1A). For eye area, we identified one significant QTL before and after accounting for body size and sex, on chromosome 3 at 3L:4.51 Mb (at genome-wide p < 0.05), explaining 3.9% of the phenotypic variance and consistent with ZI373 having larger eyes (Fig. S7A; Table 1). For inter-ocular distance, we found two significant QTL after accounting for body size and sex (at genome-wide p < 0.05), one on chromosome 2 at 2R:24.74 Mb and one on chromosome 3 at 3L:0.82 Mb, together explaining 9.3% of the phenotypic variance (Fig. S7B, Table 1). However, the effect of both of these QTL are in the opposite direction to the strain difference possibly again reflecting issues with correcting inter-ocular distance with tibia length for these strains. Tibia length variation is mostly explained by QTL on chromosome 3, at 3L:0.82 Mb, 3R:15.3 Mb (T2 tibia) and 3L:6.01 Mb (T3 tibia), and on chromosome 2, at 2L 2L:9.01 Mb (T2 Tibia) and 2L:1.6 Mb (T3 tibia) (Fig. S8A,B, Table 1).

To map intra-specific variation in eye size in *D. simulans*, we used a reciprocal backcross design, with a mapping population of 764 individuals from a cross between the Tana10 and Zom4 strains (Fig. S1B). In the Tana10 backcross, taking tibia length and sex as covariates, we identified one significant QTL on chromosome 2 at 2R:4.98 Mb, and two on chromosome 3 at 3L:5.07 Mb and 3R:16.61 Mb (all at genome-wide p < 0.05), together explaining 5.5% of the phenotypic variance in eye area, and consistent with the direction of the eye size difference between the strains (Fig. S7C, Table 2). For inter-ocular distance, taking tibia length and sex as covariates, we identified two significant QTL in the Tana10 backcross, one on chromosome 2 at 2L:9.17 Mb, and one on chromosome 3 at 3R:20.07 Mb, together explaining 2.6% of the phenotypic variance (Fig. S7D, Table 2). For the Zom4 backcross, the QTL LOD confidence intervals for eye area and inter-ocular distance variation overlap with those detected in the Tana10 backcross, with only one additional significant QTL for inter-ocular distance at 3L:16.52 Mb (Fig. S6C-E,D-F, Table 2). In both backcrosses, several QTL for tibia length were also detected above genome-wide significance (p < 0.05) on the X chromosome, and a further QTL underlying variation in the length of T2 tibia was detected at 2R:15.55 Mb in the Tana10 backcross (Fig. S8C-F, Table 2).

When comparing the mapping results for these two species, interestingly, the *D. melanogaster* and *D. simulans* maps predict a QTL for eye size at similar locations at the left end of chromosome 3, at 3L:4.51 Mb in *D. melanogaster* and at 3L:5.07 Mb or 3L:6.09 Mb in *D. simulans* (Tables 1 and 2). Although these regions encompass hundreds of genes, this finding suggests that there may be some similarities in the genetic basis of eye size variation in these two species. We found very little overlap in the positions of QTL for eye size and inter-ocular distance in either species, which may indicate that different loci underlie variation in these two traits.

### Introgression mapping of eye size differences

To verify and further refine the mapping of loci underlying variation of eye size and inter-ocular distance, we generated introgressions of either whole or partial chromosome regions between the strains used for QTL mapping. We started by reciprocally swapping each pair of chromosomes between strains except for the X, which we only introgressed from *D. melanogaster* strain ZI373 into RG13N (Fig. S2A). Not all chromosome introgressions were homozygous viable. *D. melanogaster* chromosomes 3 and 4 of ZI373 in the background of RG13N and *D. simulans*, chromosomes 2 and 4 of Tana10 in the background of Zom4 (Fig. S2C) were lethal. This is consistent with previously reported genotype ratio distortion due to genetic incompatibilities within species (CORBETT-DETIG *et al.* 2013).

In *D. melanogaster*, we confirmed that there is a significant effect of chromosome 3 on eye size where RG13N alleles are shown to decrease eye area in the ZI373 background (Fig. S6). We observed that introgression of chromosome 2 from RG13N into ZI373 also decreased the eye area suggesting that there are alleles on this chromosome that affect eye size that we did not detect in our QTL mapping (Fig. S6). However, the effect of chromosome 2 on eye area is non-reciprocal because we did not observe significant effects of the ZI373 alleles in the RG13N background when eye area residuals are examined (Fig. S6). This suggests that the effect of alleles on this chromosome on eye area may depend on epistatic interactions with loci on other chromosomes. By analyzing introgressions in the heterozygous state, we further determined that RG13N alleles generally act recessively in the ZI373 background to decrease eye size when body size is accounted for (Fig. S6A). For inter-ocular distance, chromosomes 2, and 3 from RG13N both increased this feature in a ZI373 background consistent with the strain difference although this effect was not seen with the residuals probably again reflecting an issue with correcting inter-ocular distance with tibia length (Fig. S6A).

In *D. simulans*, we found that chromosome 3 significantly affected eye area and inter-ocular distance consistent with the QTL map and direction of the strain difference (Fig. 3). Tana10 alleles on chromosome 3 act co-dominantly in the Zom4 background to increase eye area and decrease inter-ocular distance, while the Zom4 alleles on this chromosome seem to act recessively to decrease eye area and increase inter-ocular distance in the Tana10 background (Fig. 3A). When chromosome 3 was introgressed from Tana10 into Zom4 and vice versa we also observed a head shape transformation towards the strain of origin of this chromosome, suggesting that alleles on chromosome 3 are sufficient to explain a large proportion of the head shape variation between these strains (Fig. S9). Although we also detected QTL on chromosome 2 for eye area and inter-ocular distance, introgression of this chromosome from Zom4 into Tana10 only produced males with larger eyes, which is the opposite of the difference between strains (Fig. 3A).

To further investigate the position and phenotypic effects of loci on *D. simulans* chromosome 3, we generated a series of smaller introgressions with breakpoints between 1 and 5 Mb apart (Fig. S2). To estimate the position of candidate regions among these introgressions, we used a linear mixed model for eye area and inter-ocular distance variation, considering introgression line genotype as a random effect and sex and tibia length as fixed effects. We found a significant association of a proximal region on the left arm of chromosome 3 (3L:1.12-2.28Mb) with eye area (Fig. 3H,J). This region does not overlap with two other regions that have a significant effect on inter-ocular distance at 3L:10-22.1Mb and 3R:9.76-23.8Mb (Fig. 3I,J). This again indicates that different large effect loci underlie variation in these two traits. The regions identified on *D. simulans* chromosome 3 using introgressions (Fig. 3H-I) are generally consistent with the eye area and inter-ocular distance QTL positions (Fig. S7). One exception, however, was that the QTL for eye area on 3R was not recovered in the introgressions. This may have been due to epistasis because in the QTL mapping population the genetic background is recombinant with regions of heterozygosity across all chromosomes, whereas introgression lines are only recombinant across chromosome 3 while all other chromosomes are heterozygous.

### The development of eye size differences

In order to better understand the developmental basis of eye size differences between the focal strains of *D. melanogaster* and *D. simulans* used for mapping, we analyzed eye-antennal discs at 96 hours after egg laying (hAEL), when the morphogenetic furrow has moved about halfway across the presumptive retinal field. We measured the relative sizes of the eye progenitor field, as marked by expression of the retinal determinant Eya, and the differentiated part of the eye, as marked by the neuronal marker Elav (Fig. 4).

**Fig. 4.**
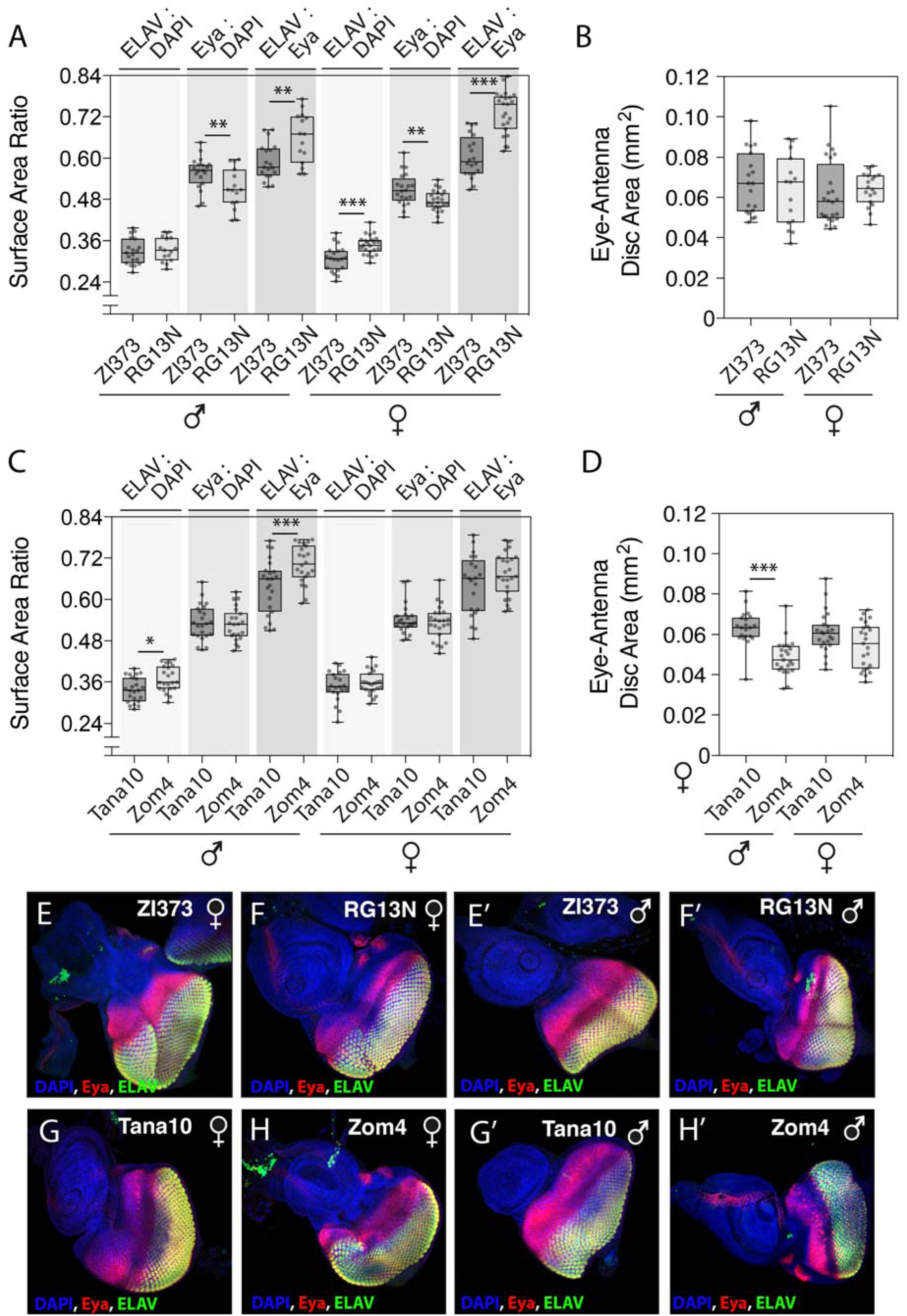
Eye fate specification at 96h AEL in *D. melanogaster* and *D. simulans*. (A,C) Ratios of the Elav, Eyes absent (Eya) and DAPI labeled regions of the eye-antenna disc and (B, D) whole eye-antenna disc areas (µm^2^) in males (♂) and females (♀) of the *D. melanogaster* strains ZI373 and RG13N and *D. simulans* strains Tana10 and Zom4, at 96h AEL. (E-H) Eye-antennal discs, dissected at 96 hAEL, of males (♂) and females (♀) of the *D. melanogaster* strains ZI373 (E, E’), RG13N (F, F’) and *D. simulans* strains Tana10 (G, G’) and Zom4 (H, H’), stained with Elav (green), Eya (red) and DAPI (blue). Scale bar = 200 µm. Statistical comparisons represent t-tests: ***p<0.0001, **p<0.001, *p<0.01.

In *D. melanogaster* there is no difference in the overall size of the eye-antennal discs among males and females of these two strains at this time point (Fig. 4B). However, the relative size of the eye progenitor field is bigger in the larger eyed strain ZI373 than in the smaller-eyed strain RG13N (Fig. 4A, E-F’). Furthermore, strain RG13N has a larger Elav-positive domain relative to the size of the whole disc in females and to the size of the Eya-positive domain in both sexes, suggestive of faster differentiation of the retina in this strain compared to ZI373 (Fig. 4A-B, E-F’). These observations suggest that eye size differences between ZI373 and RG13N may arise from both a higher number of cells committed to become eye progenitors and lower speed of retinal differentiation in the former strain compared to the latter.

In *D. simulans*, at 96 hAEL, males of the larger eyed strain Tana10 already have significantly bigger eye-antennal discs than those of Zom4, with a similar trend for females (Fig. 4D). However, we found no significant difference in the relative size of the Eya-positive domain between the Zom4 and Tana10 strains. Therefore, unlike *D. melanogaster* strains that develop big or small eyes, in *D. simulans* strains the eye progenitor field is the same size at this stage (Fig. 4C, I-L). This suggests that the difference in eye size between these *D. simulans* strains may develop later as a consequence of differences in the rate of differentiation as detected by ELAV for males (Fig. 4C) and/or the rate proliferation ahead of the morphogenetic furrow. These differences in eye development between *D. melanogaster* and *D. simulans* indicate underlying differences in the genetic basis of intra-specific variation in adult eye size between these two species.

## Discussion

### Natural variation in fly eye and head morphology

We have extended previous surveys of eye size variation within and between species of the *D. melanogaster* subgroup (NORRY *et al.* 2000; HAMMERLE and FERRUS 2003; POSNIEN *et al.* 2012; ARIF *et al.* 2013; HILBRANT *et al.* 2014; KEESEY *et al.* 2019; RAMAEKERS *et al.* 2019). Our findings emphasise the extensive natural variation in eye size among drosophilids. We further substantiate that *D. melanogaster* eyes tend to be smaller than those of its sibling species and confirm the previous finding that *D. mauritiana* tends to have the largest eyes of species in the *melanogaster* subgroup (STURTEVANT 1919; MANNING 1960; WATADA *et al.* 1986; MCNAMEE and DYTHAM 1993; POSNIEN *et al.* 2012; ARIF *et al.* 2013; HILBRANT *et al.* 2014).

We also found that there is considerable variation in ommatidia number between strains of *D. melanogaster* and *D. simulans*, with differences of up to 133 ommatidia (Fig. 2A), which could affect the vision of these flies. As suggested by our previous work, the larger eyes may have proportionally more of the ‘pale’ ommatidial subtype that detect short wavelengths of light in the UV and blue light ranges (HILBRANT *et al.* 2014), but this remains to be tested. In addition, strains with higher ommatidia number may also have higher visual acuity (CURREA *et al.* 2018; RAMAEKERS *et al.* 2019). Interestingly, however, decreasing eye size in *D. melanogaster* via nutritional restriction can result in differences of over 400 ommatidia with little detectable change in spatial acuity or contrast sensitivity because of compensation by neural summation at the expense of temporal acuity (CURREA *et al.* 2018).

We also found that flies with relatively more ommatidia have narrower ommatidia diameters and *vice versa*. This suggests that there may be a trade-off between ommatidia number and size, which may reflect packing and organization constraints in the compound eye. Changes in ommatidial diameter are also expected to have an impact on inter-ommatidial angles across the eye, and thus contribute to differences in visual abilities. Indeed, differences in inter-ommatidial angles from the center to the periphery of the eye have been described to have an important impact on spatial acuity in other dipterans, including houseflies, hoverflies and blow-flies (LAND 2009). For instance, predator flies of the genus *Conosia* display high spatial acuity and sharp gradients of inter-ommatidial angles from the center to the periphery of the eye, compared to their *Drosophila* prey (GONZALEZ-BELLIDO *et al.* 2011). Therefore in future, it will be interesting to explore if the differences in ommatidia number and diameter that we have found also result in changes to spatial acuity and contrast sensitivity, or if these are also partially compensated by neural summation, as during nutritional restriction (CURREA *et al.* 2018).

### The genetics of reciprocal changes in eye size and inter-ocular distance

Consistent with previous studies, we have also found that there is a negative correlation between eye size and other head capsule traits like face width or inter-ocular distance (NORRY *et al.* 2000; HAMMERLE and FERRUS 2003; POSNIEN *et al.* 2012; ARIF *et al.* 2013). Therefore, there appears to be a developmental trade-off between eye and head capsule size, potentially to constrain overall head size and perhaps to preserve aerodynamics. However, our mapping results for both *D. melanogaster* and *D. simulans* have revealed that there is very little overlap in the positions of QTL peaks underlying eye area and inter-ocular distance (Fig. 3, Tables 1,2). Furthermore, our higher resolution mapping of introgressions on chromosome 3L of *D. simulans* showed that different loci are responsible for variation in eye area and inter-ocular distance (Fig. 4). This is consistent with the previous observation of independent large-effect QTL underlying variation between *D. simulans* and *D. mauritiana* eye size differences, based on ommatidial diameter, and inter-ocular distance variation (ARIF *et al.* 2013). This suggests that differences in eye size caused by either changes in ommatidia number or diameter can be genetically de-coupled from the observed reciprocal change in inter-ocular distance. This further supports the notion that there is a trade-off to perhaps constrain overall head size, but may also explain why these two traits can evolve independently in some lineages of flies (GRIMALDI and FENSTER 1989; WILKINSON and REILLO 1994; SUKONTASON *et al.* 2008).

### The genetic basis of eye size may differ within and between Drosophila species

Our mapping of eye size differences between strains of *D. melanogaster* and *D. simulans* revealed overlapping QTL at the left end of chromosome 3 (Fig. 3, Tables 1,2). This suggests there could be some common genetic basis for intra-specific variation in eye size in these two species. However, we found evidence that different developmental mechanisms underlie natural variation in eye size within *D. melanogaster* and *D. simulans* (Fig. 5), which implies that different genes actually underlie the differences between these species.

It was recently shown that a SNP (*Dmel*r6.25 4:710326) in the regulatory region of *ey*, which is on chromosome 4, explains variation in eye size as a result of ommatidia number differences between *D. melanogaster* strains and between *D. melanogaster* and possibly other species, such as *D. pseudoobscura* (RAMAEKERS *et al.* 2019). Intriguingly, for this SNP, both *D. simulans* strains have the ‘A’ allele that is associated with larger eyes, while the two *D. melanogaster* strains have the ‘G’ allele consistent with smaller eyes (ALMUDI and MCGREGOR 2019; RAMAEKERS *et al.* 2019). This indicates that variation in this allele doesn’t contribute to these intra-specific differences. However, since *D. simulans* strains tend to have larger eyes than *D. melanogaster*, this suggests that this SNP in *ey* may contribute to differences in eye size between the two species, but this remains to be tested.

Finally, given previous studies and our findings that eye size differences are polygenic in *D. melanogaster and D. simulans*, with QTL on chromosomes 2 and 3, this suggests that the gene regulatory network that specifies eye size can evolve at multiple different genetic nodes. We were able to verify and narrow down the effect of one of these QTL in *D. simulans* to a relatively small region of 1.16 Mb on the left tip of chromosome 3, containing 265 annotated genes. RNA-Seq analysis of genes expressed in the eye-antennal disc at 3 developmental stages suggests that only 99 of these are expressed in this tissue (TORRES-OLIVA *et al.* 2016). Therefore, it may be possible to focus on candidate gene approaches to narrow down the genetic basis underlying differences in eye size within species. This will provide further insights into the evolution of the gene regulatory networks underlying phenotypic changes more generally, particularly those for organ shape and size (STERN and ORGOGOZO 2008; STERN and ORGOGOZO 2009; STERN 2011; MARTIN and ORGOGOZO 2013; KITTELMANN *et al.* 2018).

## Acknowledgements

This work was funded by ERC (242553) and BBSRC (BB/M020967/1) grants to A.P.M. We thank the lab of John Pool for kindly providing *D. melanogaster* strains from Zambia (ZI) and Rwanda (RG) populations and we thank the lab of Christian Schlötterer for kindly providing *D. simulans* and *D. mauritiana* strains. We thank the Cambridge Fly facility for performing embryo microinjections. We thank Linta Kuncheria for help with phenotyping of the *D. simulans* QTL mapping population and Madeleine Lindsay for help with face landmark registration. Synchrotron scans were performed at the I13-2 beamline at Diamond Light Source, UK (proposal MG23250), and we thank Kazimir Wanelik and Christoph Rau for their support. Pilot scans were performed at the TOMCAT beamline at the Swiss Light Source, Paul Scherrer Institut, Villigen, Switzerland (proposal number 20180832), and we thank Christian Schlepütz for his assistance.

---

**Fig. S1.**
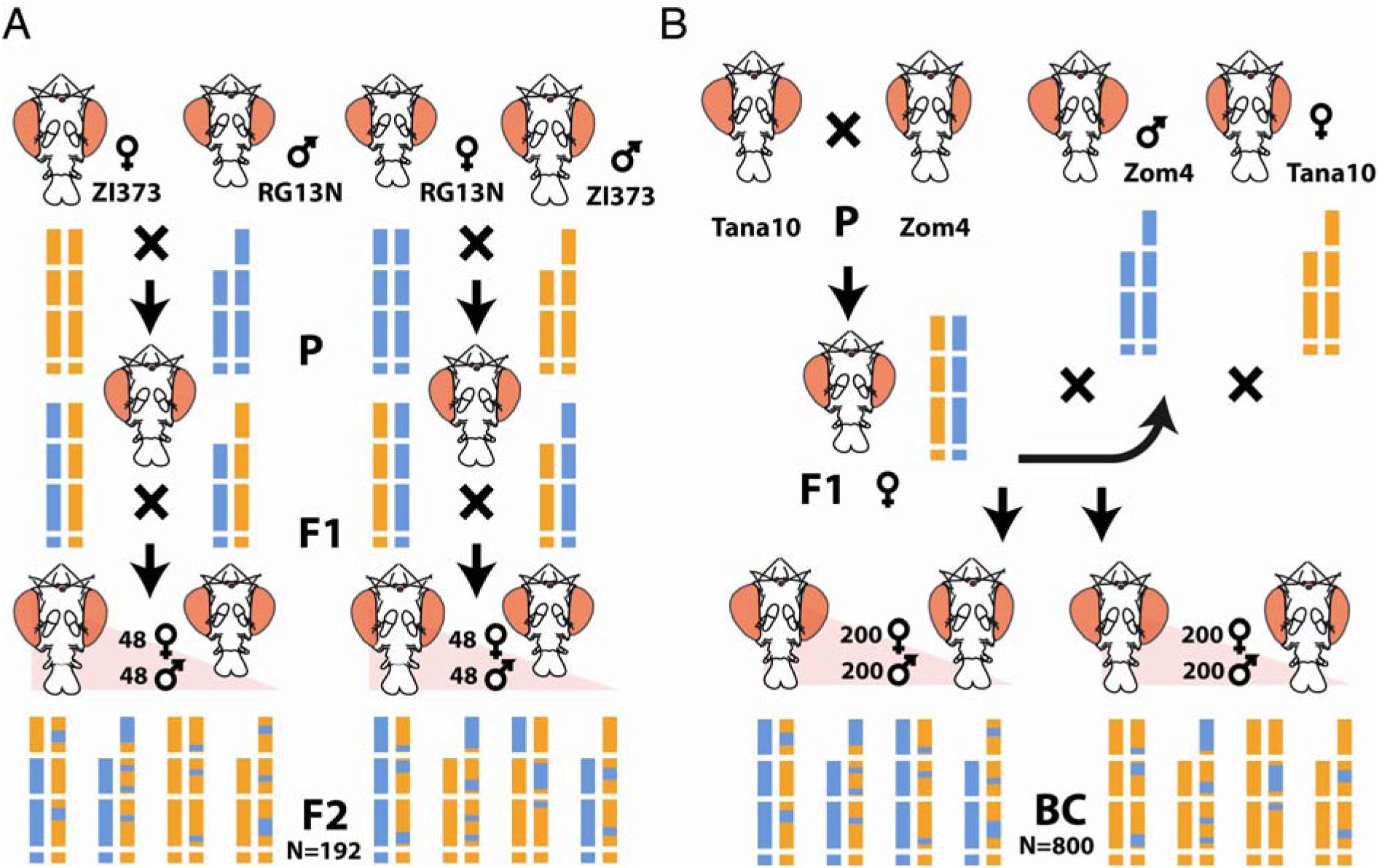
QTL mapping cross designs. (A) F2 design for crosses between the *D. melanogaster* strains ZI373 (orange) and RG13N (blue), representing the generation of 192 F2 individuals, 48 of which were female (♀) and 48 male (♂). (B) Backcross design for crosses between the *D. simulans* strains Tana10 (orange) and Zom4 (blue), representing the generation of backcross (BC) individuals, 384 from each backcross direction, each with 192 females (♀) and 192 males (♂). Separate vertical bars coloured according to expected genotypes represent the three autosomes and the X chromosome. For both mapping crosses examples of possible F2 and BC genotypes are shown.

**Fig. S2.**
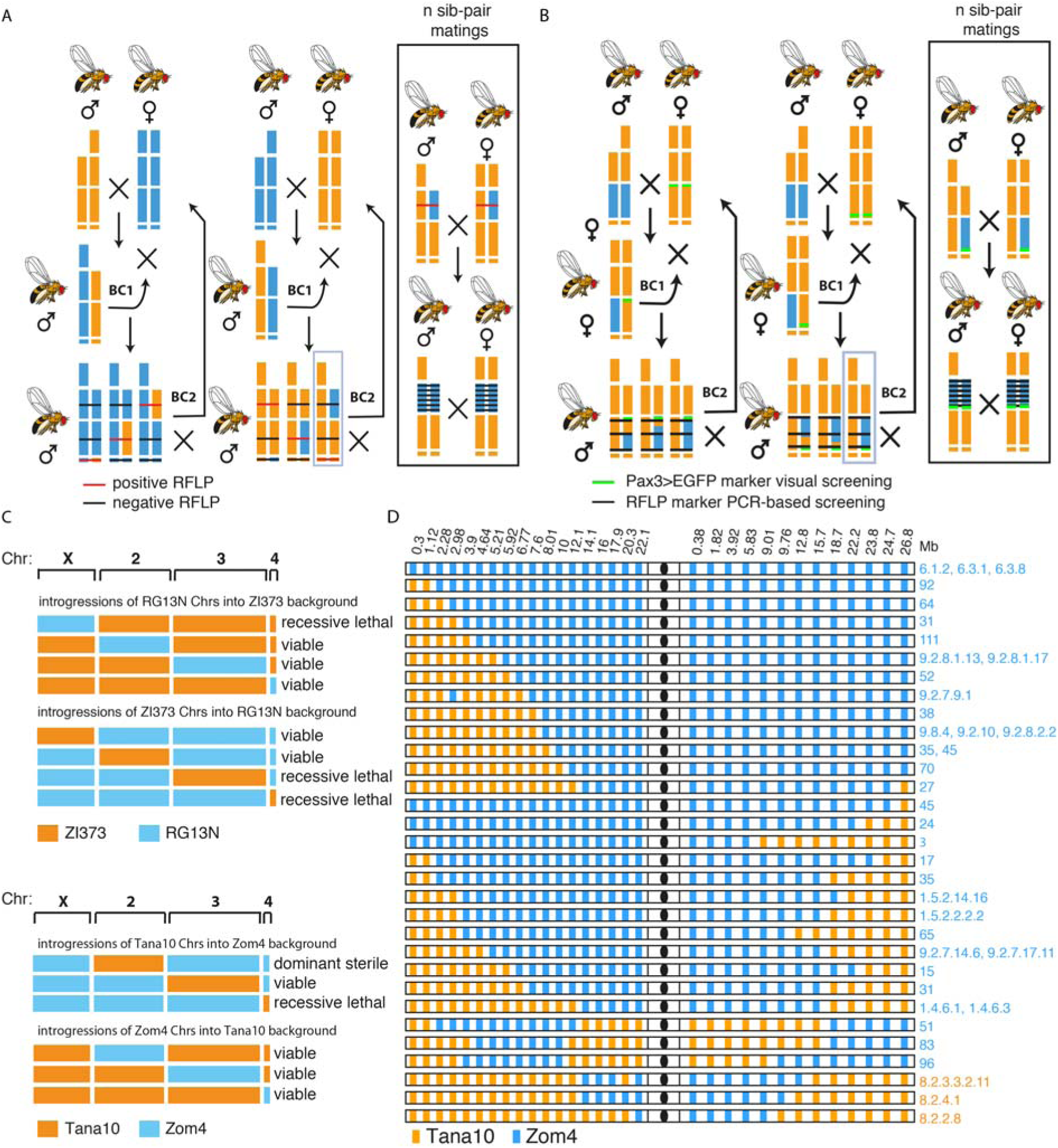
Generation method, viability and genotype summary of introgression lines. (A) Cross design used to produce large or whole chromosome introgressions between RG13N and ZI373 or between Zom4 and Tana10. (B) Cross design used to produce small 1 to 5Mb serial introgressions of Zom4 chromosome 3 (blue) in the Tana10 genetic background (orange). (C) Introgression viability summary for both *D. melanogaster* ZI373 (orange) and RG13N (blue) whole chromosomes (Chrs) or *D. simulans* Tana10 (orange) and Zom4 (blue) whole chromosomes. (D) Chromosome genotype summary all partial introgressions between Zom4 (blue) and Tana10 (orange) along RFLP genetic marker positions. The centrosome position is indicated as a black circle and genetic position is ordered in Mb from left to right starting either at the left tip/telomere side of the left arm or from the centromere side of the right arm. Apart from chromosome 3, all lines were generated by backcross to Tana10 and have a Tana10 genetic background, except for lines 8.2.3.3.2.11, 8.2.4.1 and 8.2.2.8, which were generated by backcross to Zom4.

**Fig. S3.**
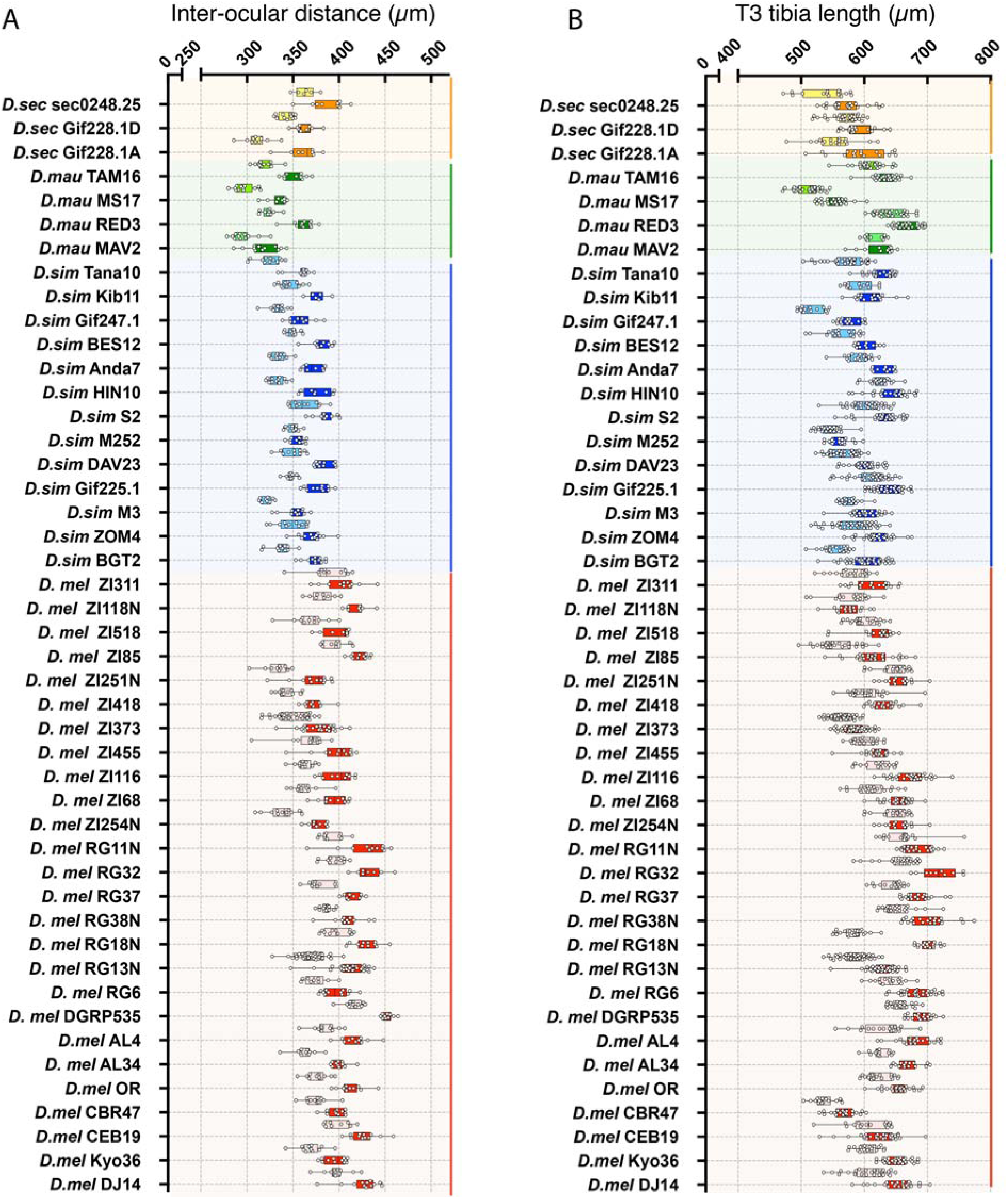
Survey of inter-ocular distance and T3 tibia length within and between species of the *D. melanogaster* species subgroup. (A,B) Box and whisker diagram of inter-ocular distance (A) and posterior tibia length (B) in strains of *D. sechellia* (males – yellow, females – orange), *D. mauritiana* (males – light green, females – dark green), *D. simulans* (males – light blue, females – dark blue) and *D. melanogaster* (males – pink, females – red) (see Table S11 for means, standard deviations and sample sizes and Table S12 for summary statistics).

**Fig. S4.**
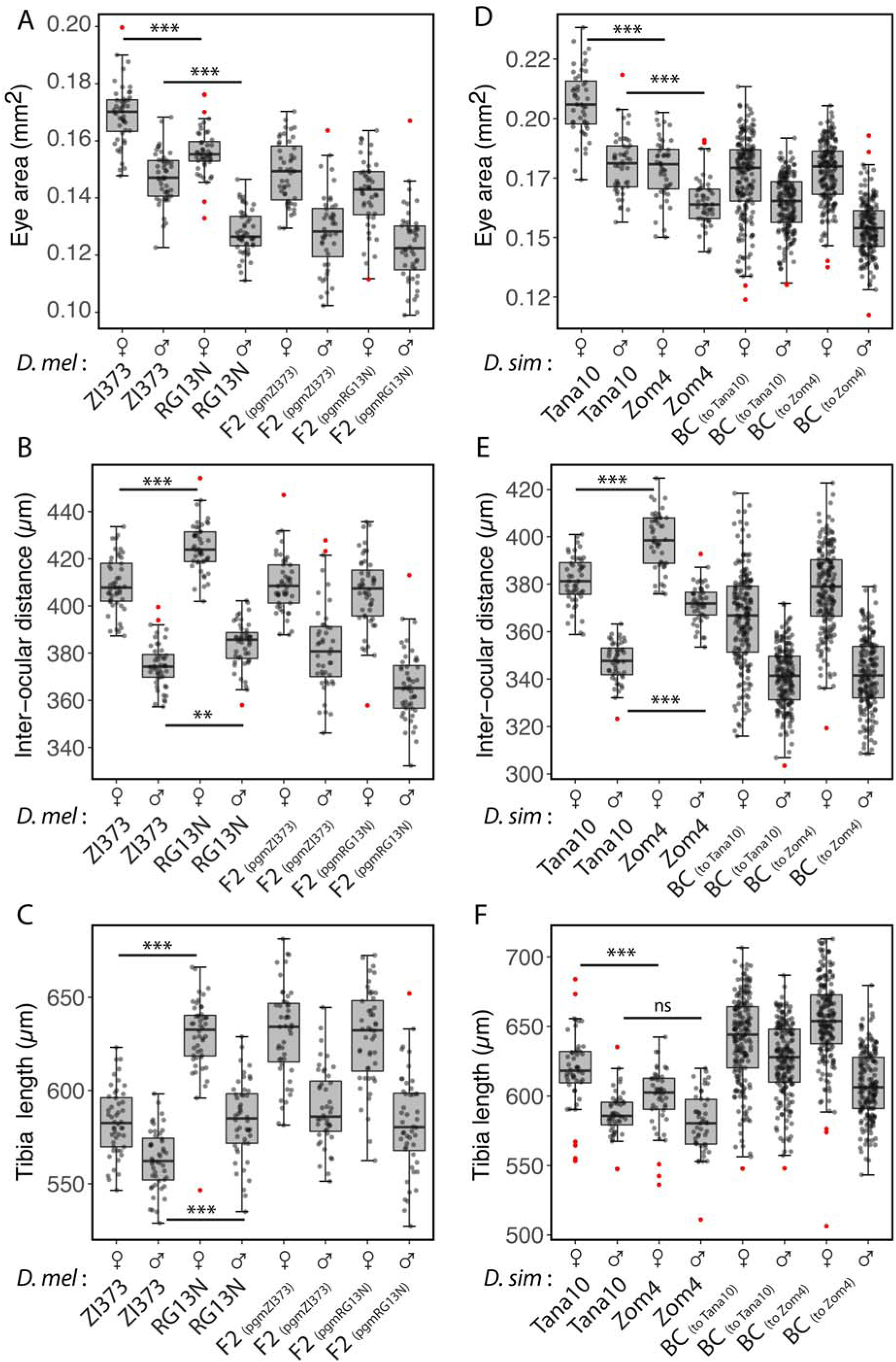
Distribution of phenotypes comparing focal strains of *D. melanogaster* and *D. simulans* and their respective QTL mapping populations. (A) Eye area, (B) inter-ocular distance and (C) posterior tibia length for females (♀, n = 45) and males (♀, n = 45) of the strains ZI373, RG13N and the F2 progeny with either parental grandmother (pgm) (n = 96×2). (D) Eye area, (E) inter-ocular distance and (F) T3 tibia length for females (♀, n = 45) and males (♂, n = 45) of the strains Tana10, Zom4 and backcross to either Tana10 (n = 192) or Zom4 (n = 192). Statistical comparisons represent pairwise t-tests: *** p<0.0001; **p<0.001, ns – p>0.01.

**Fig. S5.**
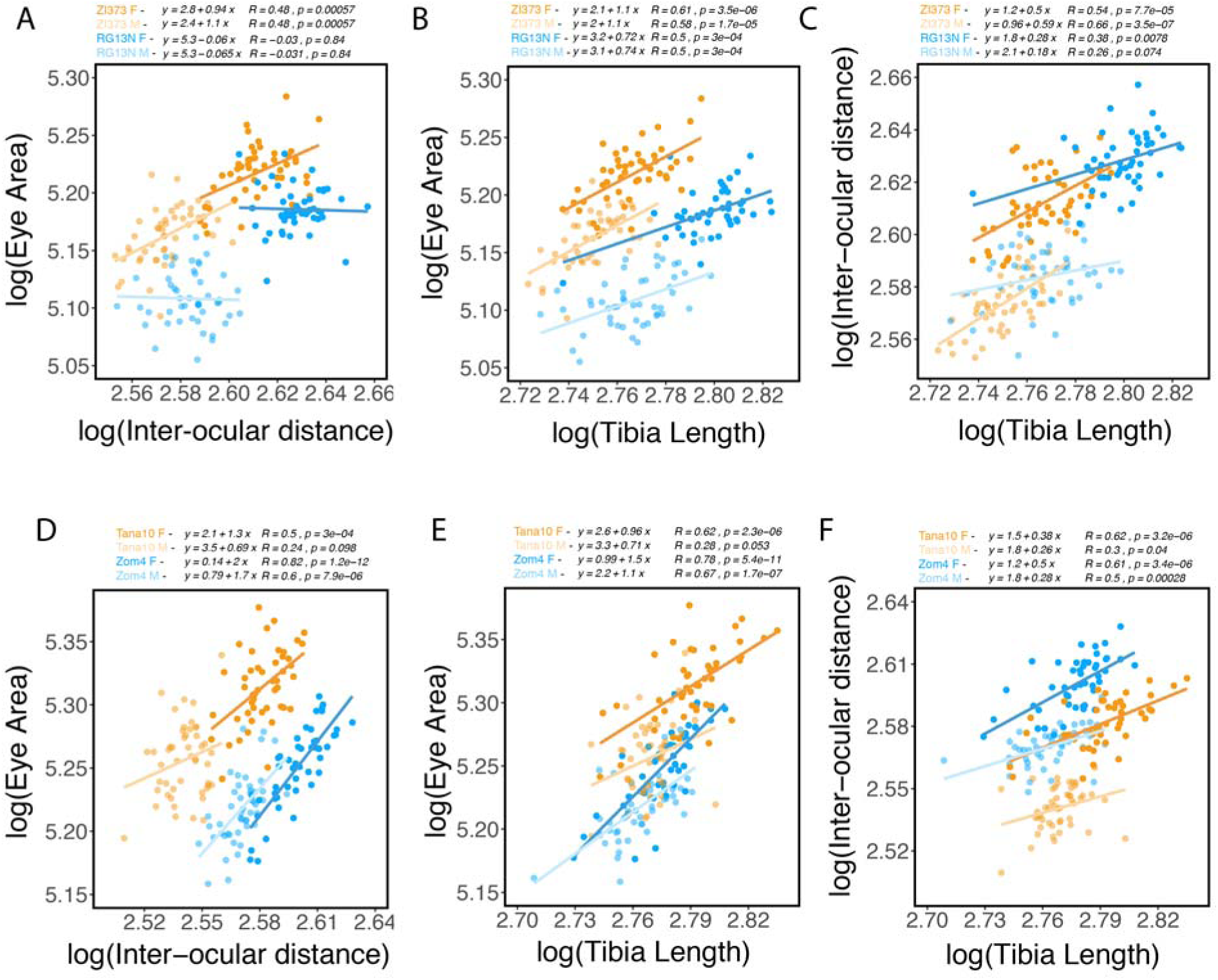
Relationships between head and body size traits. (A) Regression of eye area with inter-ocular distance, (B) eye area with the length of the posterior tibia and (C) inter-ocular distance with the length of the posterior tibia (T3), for males (M, n=45) and females (F, n=45) the strains ZI373 (orange) and RG13N (blue). (D) Regression of eye area with inter-ocular distance, (E) eye area with the length of the posterior tibia and (F) inter-ocular distance with the length of the posterior tibia (T3), for males (M, n=45) and females (F, n=45) of the strains Tana10 (orange) and Zom4 (blue). Regression equations, Pearson’s coefficient (R) and p-values are indicated above the plots.

**Fig. S6.**
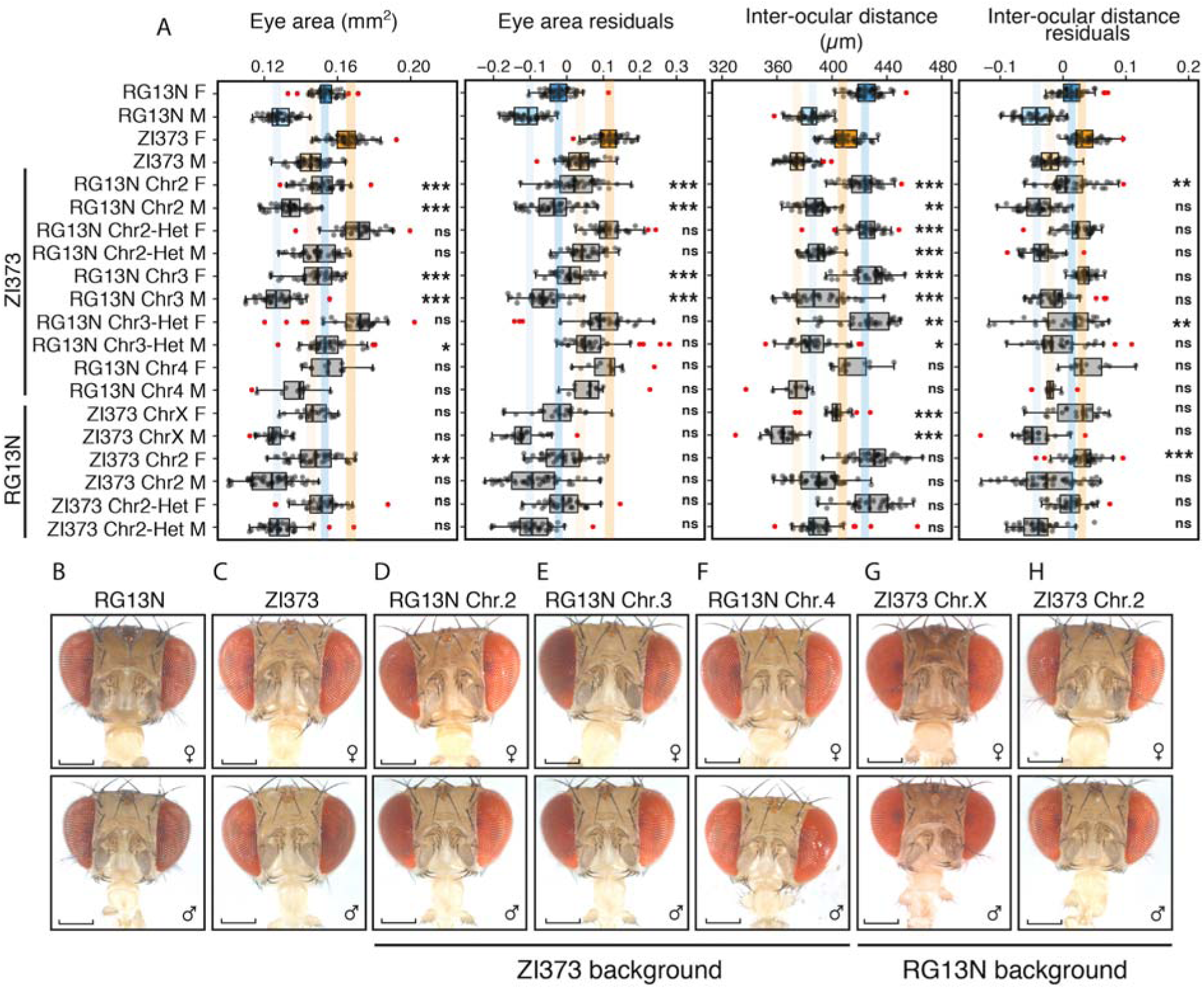
*D. melanogaster* introgressions. (A) Box and whisker diagram of eye area and inter-ocular distance, as well as the respective residuals of regression with the length of the posterior tibia (T3), measured for both females (F) and males (M) of the parental strains RG13N and ZI373 and introgression of chromosome 2 (Chr2), chromosome 3 (Chr3) and chromosome 4 (Chr4) in homozygous or heterozygous state (Het) from these strains into either genetic background. (B-G) Frontal head views of females (♀) or males (♂) from strains representative of the average of the parental strains (B) RG13N and (C) ZI373 comparing with introgression of (D) chromosome 2, (E) chromosome 3 and (F) chromosome 4 of RG13N into the ZI373 background and of (G) chromosome 2 of ZI373 into the RG13N background. scale bar – 200µm. Statistical comparisons represent pairwise t-tests by sex between introgression line and the parental strain to which this line was backcrossed to (black asterisk): ***p<0.0001, **p<0.001, *p<0.01, ns – p>0.01.

**Fig. S7.**
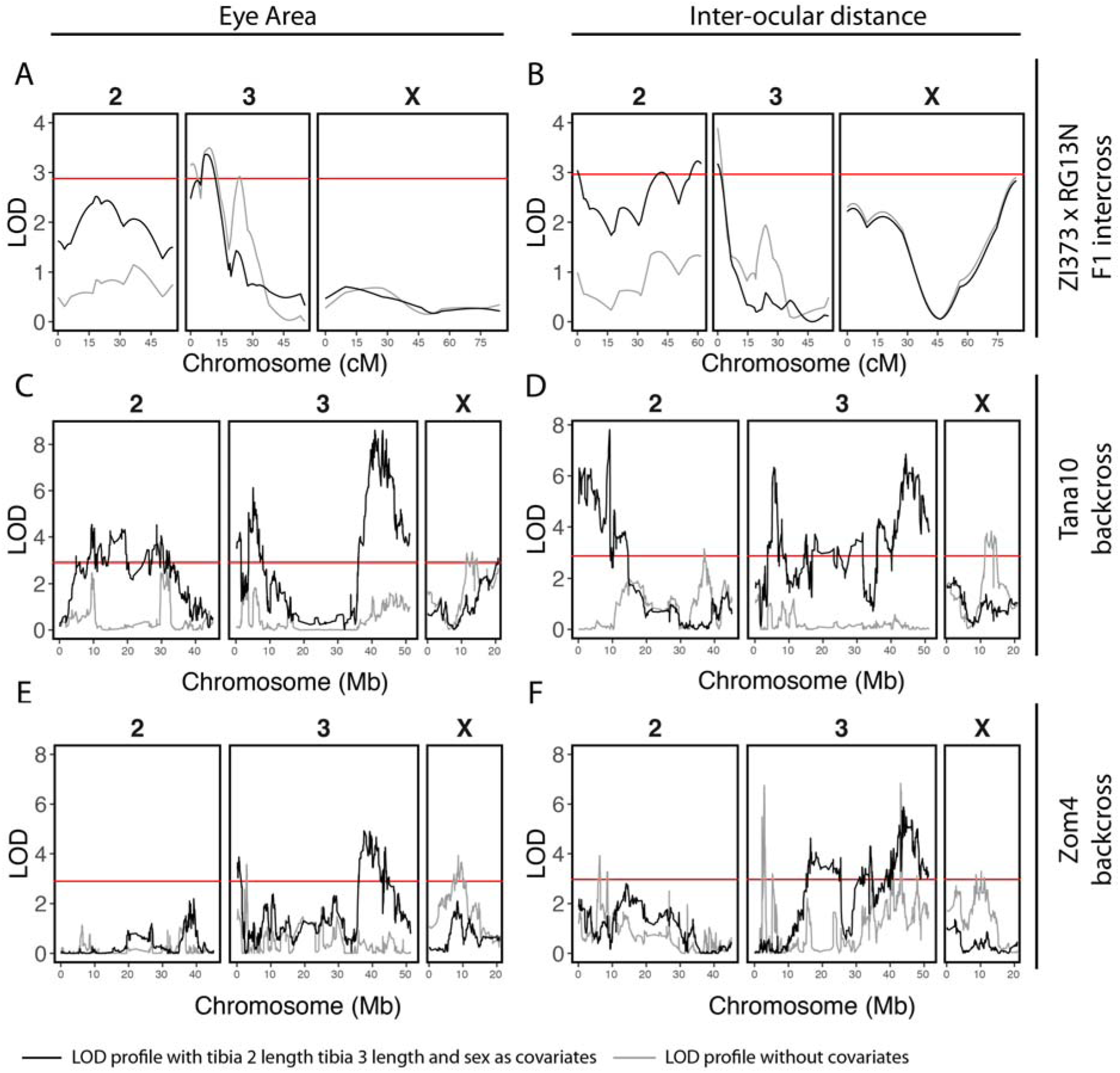
QTL maps for eye area and inter-ocular distance. (A) Eye area and (B) inter-ocular distance QTL maps based on ‘scanone’ Haley-Knott regression with *D. melanogaster* F2 progeny from reciprocal crosses between strains ZI373 and RG13N (n=192). (C) Eye area and (D) inter-ocular distance QTL maps of a *D. simulans* backcross to the strain Tana10 (n=192). (E) Eye area and (F) inter-ocular distance QTL maps based on ‘scanone’ Haley-Knott regression with *D. simulans* progeny from backcross to the strain Zom4 (n=192). A grey line represents LOD profiles without covariates and a black line indicates LOD profiles with sex and the T2 and T3 tibia lengths as covariates. A red horizontal line in each plot represents the genome-wide significance LOD threshold of p = 0.05. Note that chromosome 4 is not shown but no QTL above the genome-wide significance LOD threshold were detected on this chromosome in any of our mapping experiments.

**Fig. S8.**
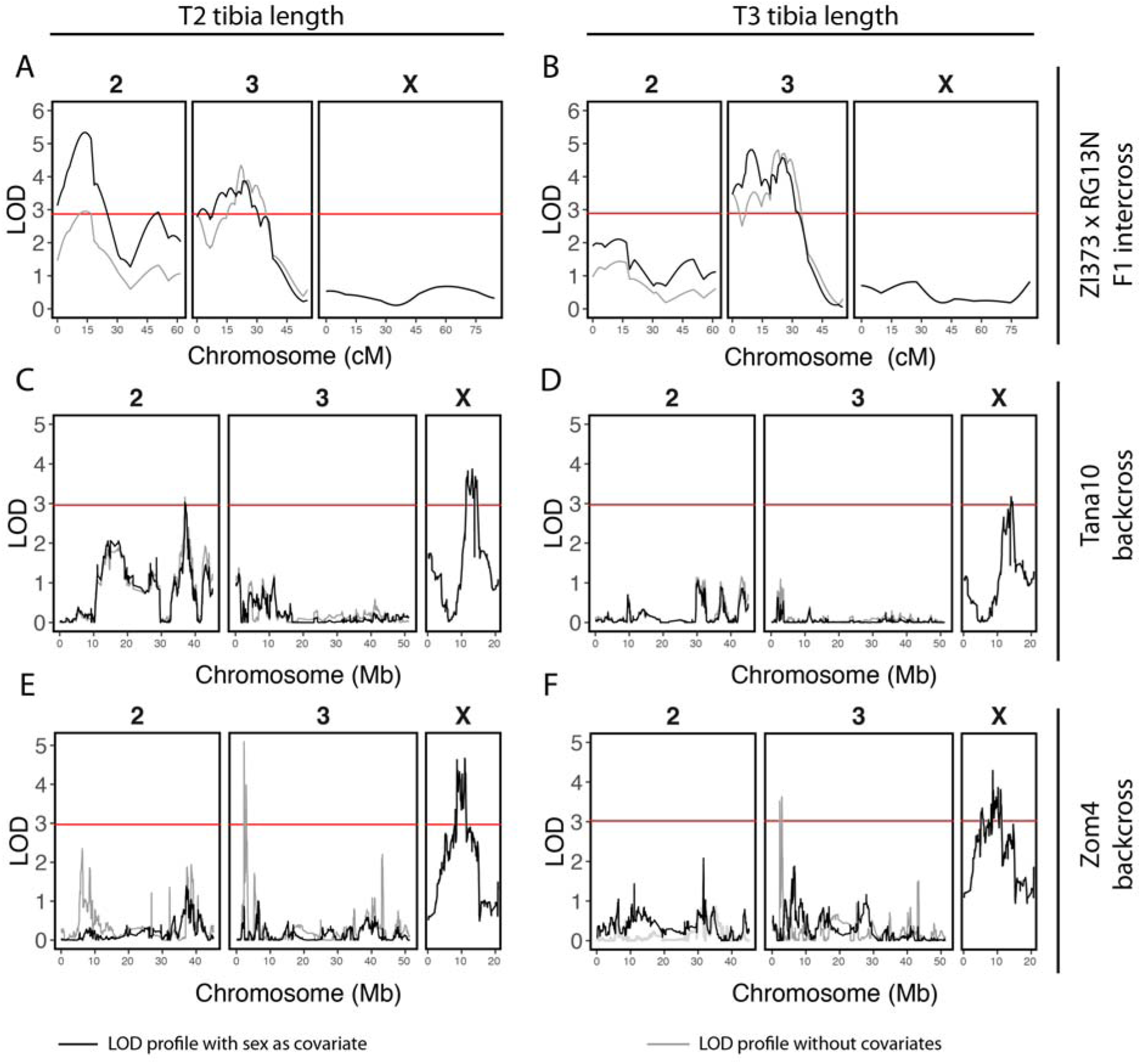
QTL maps for tibia length. (A) T2 and (B) T3 tibia length QTL maps based on ‘scanone’ Haley-Knott regression with *D. melanogaster* F2 progeny from reciprocal crosses between strains ZI373 and RG13N (n=192). (C) Middle (Tibia 2) and (D) posterior (Tibia 3) tibia length QTL maps based on ‘scanone’ Harley Knot regression with a *D. simulans* backcross to strain Tana10 (n=192). (E) T2 and (F) T3 tibia length QTL maps of a *D. simulans* backcross to the strain Zom4 (n=200). A grey line represents LOD profiles without covariates and a black line indicates LOD profiles with sex as a covariate. A red horizontal line in each plot represents the genome-wide significance LOD threshold of p = 0.05. Note that chromosome 4 is not shown but no QTL above the genome-wide significance LOD threshold were detected on this chromosome in any of our mapping experiments.

**Fig. S9.**
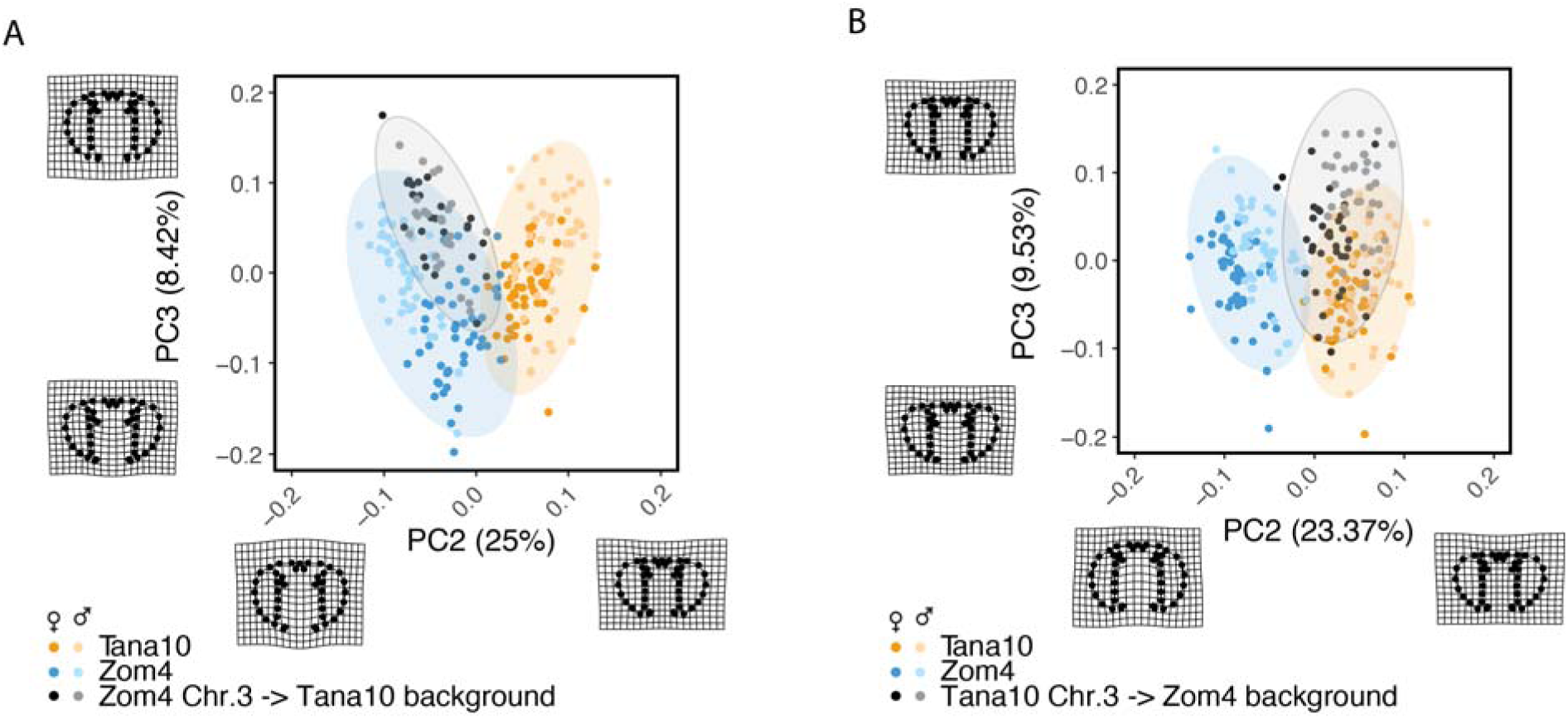
PCA of head shape variation of *D. simulans* introgression lines compared to the parental strains. (A-B) Distribution of PC2 and PC3 and their 95% confidence ellipses for position of head landmarks of the *D. simulans* strains Tana10 (orange) and Zom4 (blue) and either introgression of (A) Zom4 chromosome 3 into the Tana10 genetic background (Chr. 3) or of (B) Tana10 chromosome 3 (Chr. 3) into the Zom4 genetic background (grey). Wireframe deformation diagrams represent 2x magnifications of the minimum and maximum coordinates along the PC axis to illustrate shape differences.

